# Population structure of human-infecting and environmental isolates of *Coccidioides immitis*, the fungus causing Valley fever

**DOI:** 10.64898/2026.10.13.758980

**Authors:** Lisa I. Couper, Molly T. Radosevich, Kaylee E. Christensen, John W. Taylor, Mark C. Bitter, Katharine S. Walter, Bianca Torres, Danna Mejia, Dana Brucker, Royce H. Johnson, Justin V. Remais, Rachel B. Brem

## Abstract

*Coccidioides* is a soil-dwelling fungus that can cause serious disease (coccidioidomycosis or Valley fever) when its spores are inhaled. Limiting exposure to environmental spores is critical for disease prevention, yet how *Coccidioides* is distributed in the soil and where people become exposed remain almost entirely unknown. Here we report the first genomes of environmental isolates of *C. immitis,* from sampling of rodent burrows in the highly endemic region— California’s San Joaquin Valley. We sequenced 16 soil-derived isolates alongside 26 novel patient-derived isolates collected from the same region to investigate genetic structure and diversity in *C. immitis*, and the potential origins of human infection. We found evidence of regional population structure among environmental *C. immitis* strains, though with extensive allele sharing between clusters. Most patient isolates derived nearly 100% of their ancestry from a genetic cluster shared with soil isolates, suggesting that the *Coccidioides* strain inhaled by a human host represents a random draw from the environmental pathogen pool. Further, we found that high levels of genomic diversity in environmental *C. immitis* are partitioned into effectively clonal lineages within individual rodent burrows, with isolates from neighboring burrows differing by over 13,000 SNPs. Together, these data support a picture of diverse, regionally structured environmental populations of *C. immitis*, suggesting that strategies to assign patient exposures to general regions within the San Joaquin Valley may be possible.

## Introduction

*Coccidioides* is an emerging fungal pathogen that can cause severe disease in otherwise healthy individuals^1,2^. The fungus typically resides in the soil, often in close association with small burrowing mammals^3–5^, but can become aerosolized when soils are disturbed through processes such as windstorms, construction, or excavation^6^. Inhaling airborne spores results in the disease coccidioidomycosis (Valley fever), of which over 200,000 cases are estimated to occur each year in the U.S. alone, predominantly in the southwest^7^. There are currently no vaccines approved for human use, and limited treatment options^8–11^ as well as rising concern of antifungal resistance^12,13^. As such, the primary strategy for coccidioidomycosis prevention and control is to minimize exposure to environmental sources of the pathogen. Yet this is challenged by a lack of understanding of where the pathogen persists in the environment, how it spreads, and where human exposures occur.

*Coccidioides* spp. live in soils of arid and semi-arid regions of the Americas, with *C. immitis* primarily found in California, and *C. posadasii* found more broadly across the southwestern U.S., Mexico, and Central and South America^14,15^. Pathogen dispersal has been proposed to occur via wind, land disturbance, and human and animal movement^16–18^. Support for wind-mediated long-distance dispersal has emerged from notable outbreaks of coccidioidomycosis occurring outside the endemic region following major wind events^19,20^. A role for human migration and movement of animals in long-distance dispersal is suggested by *Coccidioides* hotspots in non-endemic regions such as Washington state and northern Utah, which are associated with ancient burial sites^17,21^. At finer spatial scales, the movement of infected mammalian hosts such as kangaroo rats and pocket mice^22,23^, whose range sizes are typically limited to the area directly surrounding their burrows, could also disperse *Coccidioides*. However, much of this understanding has been inferred from outbreaks of coccidioidomycosis, which are inherently sporadic^24^ and influenced by human risk factors and behavior^6^, while the routine ecological processes of pathogen dispersal and gene flow remain poorly understood.

For other pathogenic taxa, landscape genomic approaches—which leverage genomic variation in samples collected across space—have provided valuable insights into pathogen transmission and spread^25,26^. Namely, by comparing genetic sequences of the pathogen from infected patients to that of environmental samples, these approaches can help identify environmental sources to target for control or avoidance^26–28^. A major challenge in applying such approaches for *Coccidioides* is that environmental genomes of the pathogen have proven difficult to generate. Doing so requires extensive environmental surveillance (as the pathogen is diffuse within soils), molecular detection, and isolation, involving either costly target enrichment or *Coccidioides* culture within high-containment facilities^29–31^. Genomes generated from infected patient tissue are more readily available, but the location where an infected patient was exposed is typically unknown, limiting the spatial resolution of potential analyses on pathogen dispersal and population structure.

In this work, we draw upon nearly a decade of soil surveillance in the San Joaquin Valley of California, to develop cultures of environmental isolates of *C. immitis* from this highly endemic region. These 16 soil-derived strains were isolated from precisely georeferenced rodent burrows, enabling investigation of *Coccidioides* diversity and structure at much finer spatial resolutions (i.e., sub-kilometer) than previously possible. We apply whole genome sequencing on these soil-derived isolates alongside 26 novel patient-derived isolates from similar geographies. We ask: (i) How genetically diverse is *Coccidioides immitis* in this endemic region and how is this diversity structured across space? And (ii) do strains infecting patients reflect random draws from the environmental pool or a distinct subset of it?

## Results

### Genomic dataset of soil- and patient-derived isolates

We isolated a total of 16 soil-derived (15 collected from 2018–2022, 1 from 2025) and 26 patient-derived (all from 2024–2025) *Coccidioides immitis* from the San Joaquin Valley of California (Figure 1, Supplementary Table S1). Each soil-derived isolate originated from a rodent burrow sampled at one of five collection sites spanning this endemic region: Bakersfield, the Carrizo Plain National Monument (‘Carrizo’), McKittrick, Coalinga, or Tracy. These sites were sampled as part of the current study or prior studies by our group^3,32–34^. Within these sites, several isolates originated from a single rodent burrow, enabling investigation of genomic variation across levels of within-burrow, within-site, and between-site. The 16 soil-derived isolates were the result of culturing 237 qPCR-positive soil samples (i.e., 6.8% culture success). We found no discernible variation in culture success based on the year or month of soil sample collection (Supplementary Figure S1). All patient-derived isolates were from patients of Kern Medical in Bakersfield. Two of the latter (VFI_5 and VFI_25) originated from the same patient and were collected at the same time but from distinct within-host sites (i.e., tissue versus vertebra); all others (n = 24) were from unique patients.

**Figure 1.**
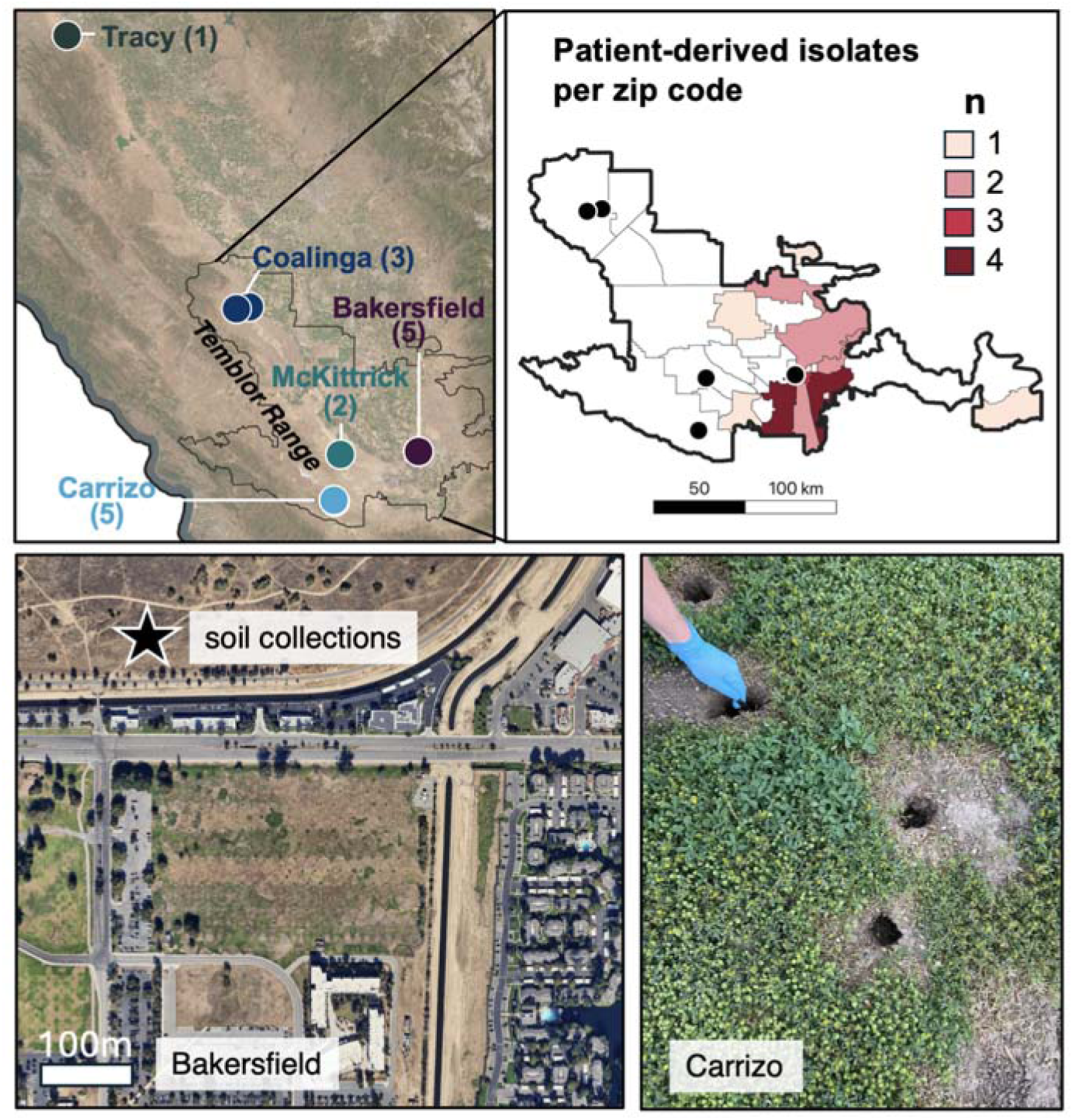
Geographic origins of *Coccidioides* isolates sequenced in this study. Geographic origin and count of our 16 soil-derived isolates (top left) or 26 patient-derived isolates (top right). Patient-derived isolates are shown at the zip code level of the patient’s home residence. Black points in the top right panel denote the location of the soil-derived isolates from Bakersfield, Carrizo, Coalinga, and McKittrick. Soil samples from Bakersfield were collected from ∼200 m along the Kern River Parkway, near an urban area (bottom left). All soil-derived isolates were collected from rodent burrows (bottom right).

We sequenced all isolates and generated genomes meeting quality control criteria, including >10x depth for at least 90% of the genome (median genome-wide depth = 178x; Supplementary Table S2). An additional 44 previously published (“legacy”) patient isolates met these criteria and were included when building the genome-wide phylogenetic tree. After joint genotyping and variant filtering, we identified 55,874 single nucleotide polymorphisms (SNPs) segregating across all samples (n = 86) and 56,282 SNPs across the novel soil- and patient-derived samples (n = 42) (after filtering SNPs with minor allele frequency < 0.05). Given a *Coccidioides* genome size of 29 Mb across six large scaffolds, of which 23.6 Mb was callable, these SNP counts constitute ∼0.24% of callable bases.

### Soil-derived genomes cluster based on regional geography and do not constitute distinct populations from patient-derived genomes

As a first-pass investigation of relatedness between our *C. immitis* isolates, we used their genomes as input into assessments of genome-wide population structure using principal component analysis (PCA) and the ADMIXTURE software, running the latter with a fixed number of population clusters (*K*) as a user-specified input from 2 to 10, and using one representative isolate per clonal set. For soil-derived samples, we found evidence of population structure corresponding to regional geography. Namely, both the PCA and ADMIXTURE results showed that soil *C. immitis* isolates clustered based on collection site, with primary separation between Bakersfield and non-Bakersfield sites at *K* = 2 and 3, and further separation of the non-Bakersfield cluster into the southwestern Central Valley (Carrizo and McKittrick collection sites), and the western Central Valley (Coalinga and Tracy collection sites) at *K* = 4 and beyond (Figure 2, Supplementary Figures S2–S4, Supplementary Results). Among soil samples, genome-wide differentiation between groups was moderate, with a weighted F_ST_ of 0.15 and 0.18–0.23 under models of two and three soil clusters, respectively, inferred by ADMIXTURE (Supplementary Table S3).

**Figure 2.**
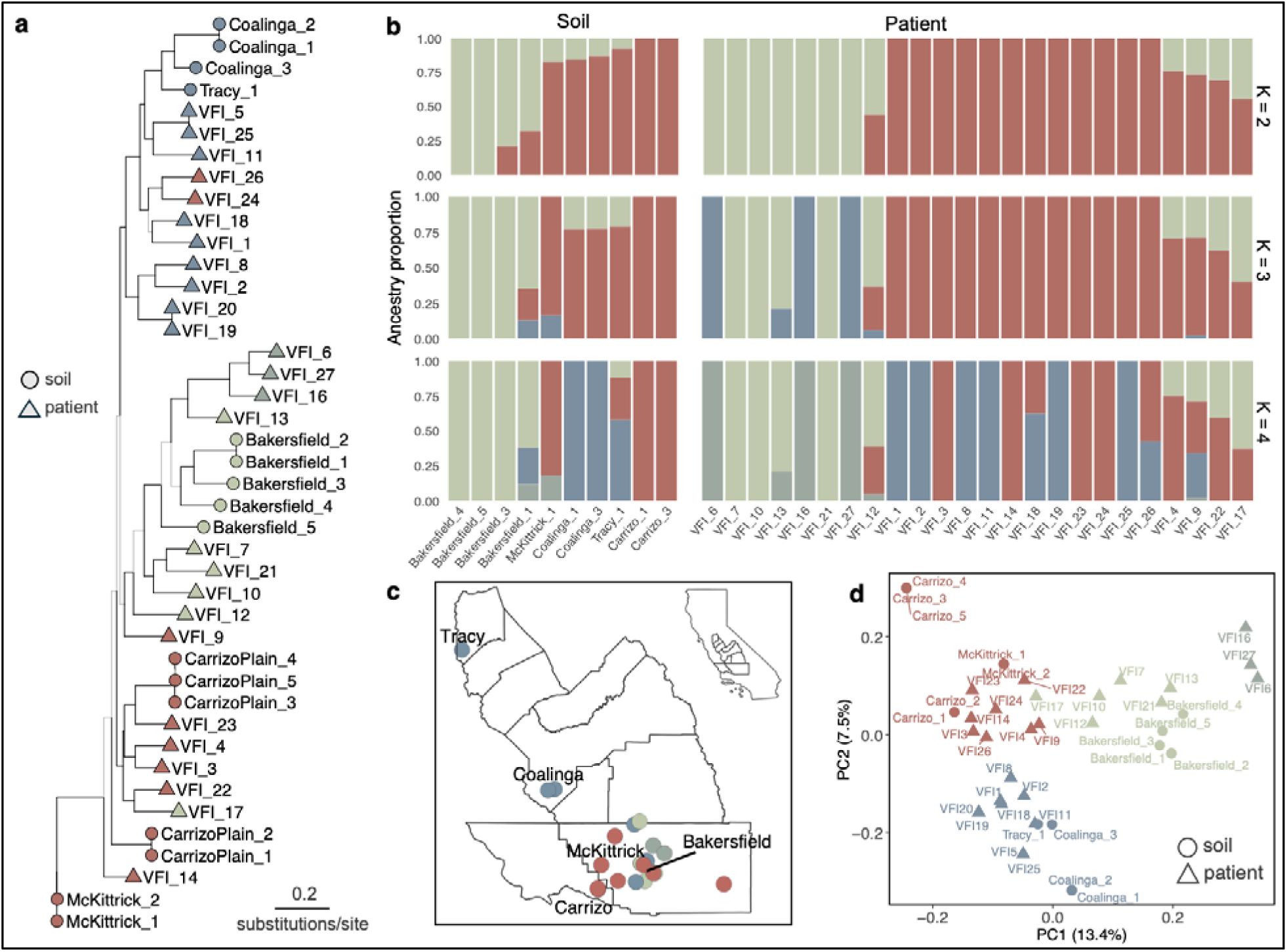
Phylogenetic relationships and population structure of *C. immitis*. **(a)** Maximum likelihood phylogeny on our *C. immitis* genomes derived from soils (circles; n = 16) or patients (triangles, n = 26). See Supplementary Figure S7 for the tree including legacy patient genomes. Gray branches have low support (SH-aLRT < 80 and/or UFBoot < 95). Tips are colored based on their dominant assignment from ADMIXTURE under *K* = 4. The tree is unrooted and branch lengths are proportional to the number of substitutions per site. **(b)** Ancestry proportions estimated from ADMIXTURE under *K* = 2–4. Each bar represents an isolate, with soil-derived on the left and patient-derived on the right. For any clonal sets, we included only one isolate in the ADMIXTURE run to avoid violating assumptions of independence, leaving 10 soil-derived and 24 patient-derived genomes. **(c)** Geographic origin of isolates (i.e., either the exact location or the zip code centroid for soil-derived and patient-derived isolates, respectively). Isolates are colored by their dominant cluster assignment under *K* = 4. **(d)** PCA of genetic variation withsoil-derived isolates (circles) and patient-derived isolates (triangles) colored by their dominant cluster assignment under *K* = 4. The first two axes are shown with the percent variance explained in parentheses. ‘VFI’ stands for the Valley Fever Institute.

Over 60% of the patient-derived *C. immitis* genomes shared full ancestry with a genetic cluster containing soil isolates, as estimated by ADMIXTURE at *K* = 2 or 3 (Figure 2B), and patient-derived *C. immitis* genomes were scattered amongst the soil isolates on the PCA rather than forming a single distinct cluster (Figure 2D). Along these lines, we found that soil- and patient-derived genomes did not constitute distinct populations, as overall genomic differentiation was minimal (F_ST_ of 0.013, permutation p = 0.14), and we detected no overall clustering based on sample source (i.e., soil versus patient) (Figure 2D, Supplementary Figure S5) or year of collection (2018–2025) (Supplementary Figure S6). The presence of environmental population structure and the lack of clear, overall differentiation between soil and patient genomes suggest that strategies to assign human exposures to geographic regions within the San Joaquin Valley may be possible.

### Shallow phylogeographic structure

As a complementary approach to investigating relationships among our strains, we constructed maximum likelihood phylogenies using the full genome for our isolates, as well as a version including legacy patient genomes from across the western U.S. and Mexico. The resulting tree indicated shallow phylogeographic structure in *C. immitis* (Figure 2A, Supplementary Figure S7). Amongst the soil *C. immitis* isolates, there was clear phylogenetic separation between those from the Bakersfield region and all other collection locations, as well as further separation between those from Carrizo and McKittrick versus Coalinga and Tracy, all with well-supported branches (SH-aLRT = 100 and UFBoot = 100). Notably, these groupings are similar to those independently inferred by ADMIXTURE, providing support for a model of regional population structure among environmental strains of *C. immitis*.

Within this broader separation, soil-derived genomes from different collection locations did not form reciprocally monophyletic groups, and genomes from soils, patients, and legacy patients were interspersed throughout much of the tree. These findings are consistent with a model of shallow divergence defining clades of *C. immitis*, with additional contributions from incomplete lineage sorting and recombination. Along these lines, we found low absolute as well as linkage disequilibrium that decayed with genomic distance (Supplementary Table S4, Figure S8), providing additional evidence of recombination in the history of our *C. immitis* populations.

The most deeply diverged lineage consisted of three patient isolates—VFI_6, VFI_16, and VFI_27. These isolates had pairwise genetic distances of ∼8,400–12,000 SNPs (Supplementary Figure S9) but shared two deletions (∼47 kb in total) with nearly identical breakpoints. These deletions were not present in any other isolate or the outgroup, suggesting they were derived (Supplementary Figure S10). Based on the annotated *C. immitis* genome (*C. immitis* RS GCF_000149335.2), the deleted region encompasses six genes, each of which is currently categorized as hypothetical protein. These data support an assignment of VFI_6, VFI_16, and VFI_27 as a clade whose ancestry is in part attributable to sources unsampled in our current set. These three strains were isolated from distinct patients that did not share a zip code or other discernible metadata (e.g., disease manifestation, body site of collection).

### Soil isolates are clonal within a burrow, but divergent across burrows at any distance

To understand the spatial extent of dispersal in *Coccidioides*, we investigated patterns of genomic distance across scales of within-burrow, between-burrow (but within a collection site), and between collection sites. We found that *C. immitis* isolates from the same burrow were clonal or near-clonal. That is, pairs of soil isolates originating from the same burrow differed by 29–169 SNPs (Figure 3). This was on par with three technical replicates in which a given isolate was re-cultured and re-sequenced, and differed from its original isolate by 47–80 SNPs (Supplementary Results, Supplementary Figure S11). Given the finding of within-burrow clonality, we retained only one isolate per burrow in population structure and downstream analyses (n = 10; described further below) to avoid violating assumptions of independence. A complementary set of results using all samples is given in the Supplement.

**Figure 3.**
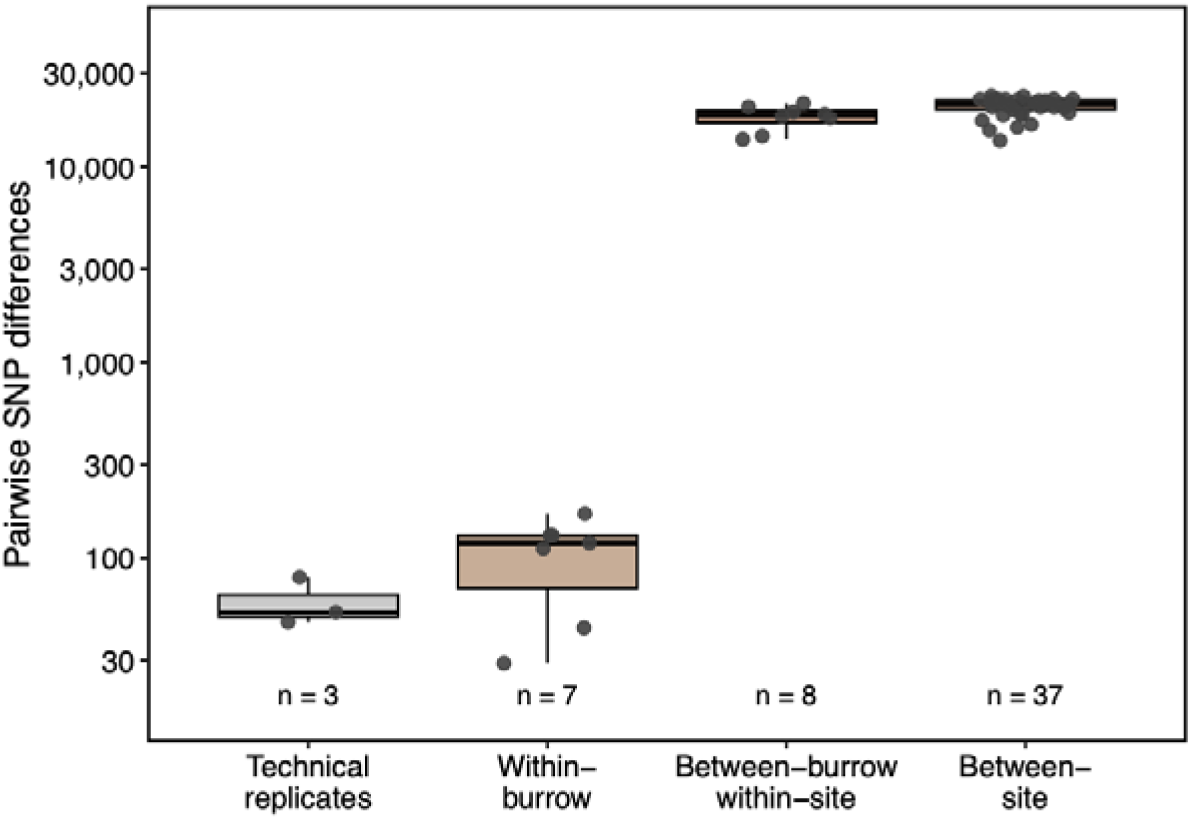
Genetic distance between soil-derived *C. immitis* isolates. Box plot of pairwise SNP differences across scales of within-burrow, between-burrow within the same collection site, and between collection sites. Pairwise differences between technical replicates (the same isolate re-cultured, extracted, and sequenced) are included for comparison. Note that the y-axis is on a log-10 scale to aid visualization of differences between groups.

Each soil *C. immitis* isolate harbored an average of 3,279 SNPs not found outside of its burrow (range: 2,214–4,551), and 52.3% of all SNPs segregating among the soil samples were private to a single burrow, indicating that the lineage at a given burrow harbored substantial unique variation (Supplementary Table S5). A rarefaction curve using data from isolates from across the ten burrows showed no apparent saturation in genomic variation (Supplementary Figure S12). The asymptote of this curve, which represents a lower bound estimate of the total SNPs segregating across our soil populations (i.e., the Chao2 estimator), was 110,200 SNPs, of which we captured 62,759 (57%), suggesting that many rare variants in environmental *C. immitis* remain undetected.

Beyond clonality within a given rodent burrow, the extent of physical separation between burrows had no further detectable impact on divergence levels of the *C. immitis* strains from those burrows (Figure 3). Specifically, soil isolates from different burrows at the same site (typically <1 km apart) differed by 13,811–21,111 SNPs (mean: 17,898), whereas soil isolates collected from distinct sampling sites—separated by tens to hundreds of kilometers—differed by 13,627–23,126 SNPs (mean: 20,300) (Figure 3, Supplementary Table S6). Further, we detected no significant scaling between the mean genomic distance between isolates from distinct sampling sites and the geographic distance between those sites (R^2^ = 0.009, p = 0.80) (Supplementary Figure S13, Table S6), although our power to detect isolation-by-distance was limited (i.e., 18% power at the observed effect size; Supplementary Results). Thus, at both the between-burrow scale and the between-site scale, we found no evidence that geographic separation predicted genomic separation. This is consistent with our population inference results overlaid on the geographic origin of isolates, wherein, for example, soil isolates from Tracy and Coalinga (197 km apart) were assigned to the same population (Figure 2C). We interpret these results as indicating that *C. immitis* dispersal can be extensive at the regional scale, such that allele-sharing can occur between strains isolated from very close or distant locations; furthermore, the near-clonality of isolates from within a given burrow reflects the maintenance of a given genotype deposited there.

### Soil- and patient-derived genomes are similarly diverse

We next compared patterns of genomic diversity among patient and soil *C. immitis* isolates. We considered that, in principle, patient isolate diversity could be lower than that of isolates from soils if human infectivity were associated with a strong bottleneck for *Coccidioides*. We calculated several measures of genomic diversity including Watterson’s estimator and nucleotide diversity among the patient and soil isolates (n = 24 and 10, respectively, after removing near-clones), finding that each of these measures was broadly similar between the groups (Supplementary Table S7). Namely, nucleotide diversity among the patient isolates was not lower than that of the soils (6.58 x 10^-4^ vs 6.34 x 10^-4^, respectively). Thus, our data afforded no evidence for a genetic bottleneck among clinical *C. immitis*, to the extent detectable from our sample size.

We further explored potential drivers of genetic diversity by examining the distribution of mating type loci idiomorphs (*MAT1-1* and *MAT1-2*) across our *C. immitis* strain genomes, given that sexual reproduction can only occur between isolates of differing idiomorphs. Both idiomorphs were present among the pool of soil-derived isolates, and, separately among the patient-derived isolates, at similar frequencies (Supplementary Table S8; Fisher’s exact test odds ratio: 2.68, p = 0.27). For soil isolates specifically, those originating from the same burrow (5 sets of 2–3 isolates) always shared the same idiomorph (Supplementary Table S8), consistent with a model of clonal growth within the burrow.

### Most patient isolates share ancestry with soil isolates

We hypothesized that soil *C. immitis* isolates could represent source populations from which genome segments in clinical isolates were inherited. To test this, we used *Twisst* (topology weighting by iterative sampling of subtrees)^35^ to estimate the support for different phylogenetic relationships at windows along the genome, using the soil *C. immitis* isolates as reference groups. For each genomic window, the *Twisst* method infers support for a topology in which a focal isolate is either in a clade sister to one of the soil reference groups or on its own branch, the latter reflecting either a divergent lineage or incomplete lineage sorting and estimation error. As a calibration, we noted that the data most strongly supported the placement of a given soil-derived isolate in a clade with a soil reference population in 73.4% of its genomic windows, on average (Figure 4B, Supplementary Table S9). Patient-derived isolates were nearly indistinguishable from this baseline, as the placement of a given patient-derived strain with a soil reference population was the most strongly supported topology across 72.7% of its genomic windows, on average. The latter result excludes three patient isolates—VFI_6, VFI_16, and VFI_27—the same isolates that formed a divergent lineage according to the phylogenetic tree (Figure 2A). Each of these isolates was more commonly placed on its own branch, doing so across 49.3% of windows, on average, indicating that the *Twisst* approach can detect divergent ancestry when present. Thus, our results suggest that patient-derived isolates more commonly share ancestry with sampled soil populations rather than constituting divergent lineages, with the exception of VFI_6, VFI_16, and VFI_27.

**Figure 4.**
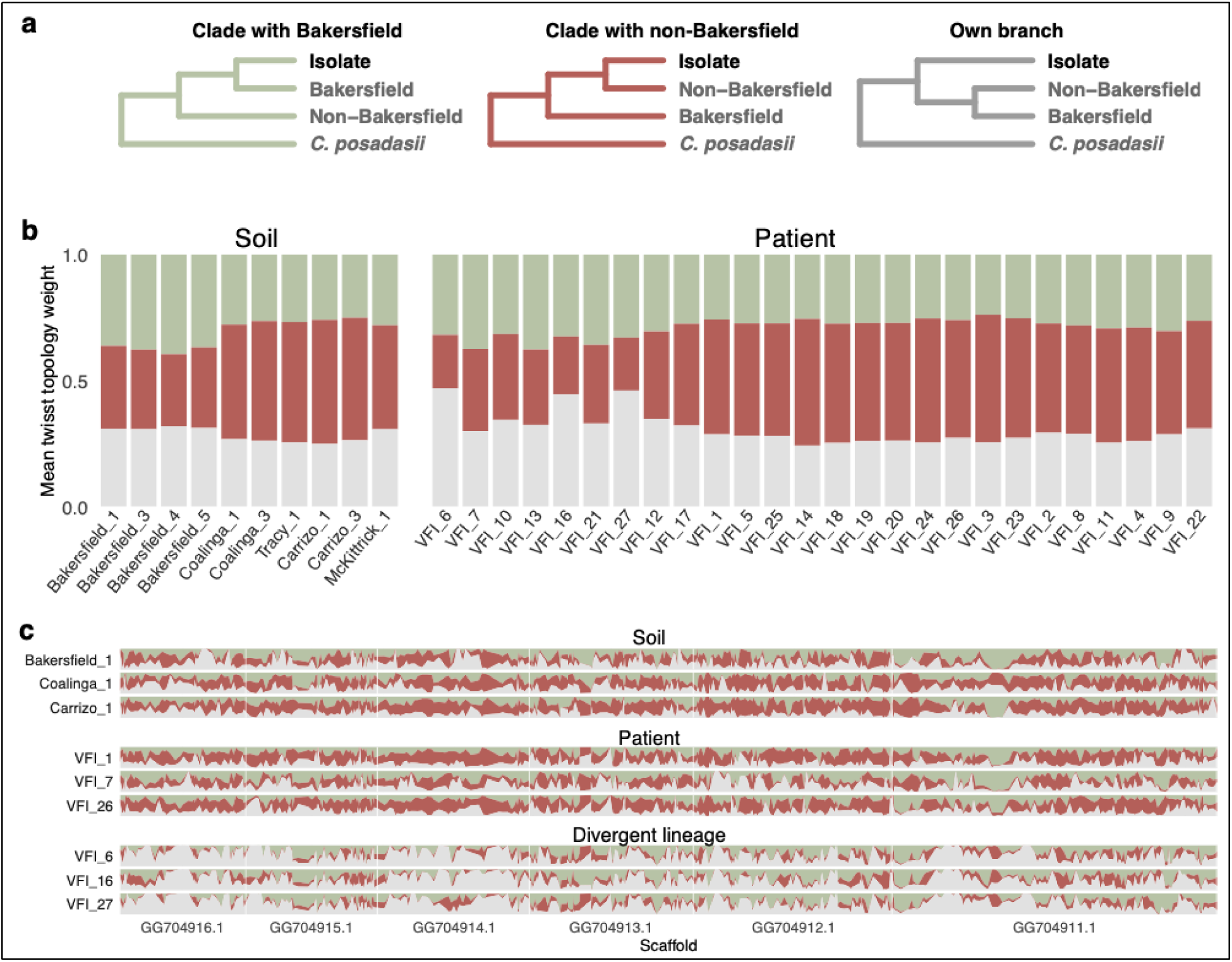
Phylogenetic relationships among soil and patient-derived isolates. **(a)** The three possible rooted topologies given two populations, one focal isolate, and an outgroup (*C. posadasii*). **(b)** Weightings for each topology, averaged across the genome, for each soil (left) or patient (right) isolate. Note that VFI_5 and VFI_25 are *C. immitis* isolates from the same patient; all others are from distinct patients. **(c)** Weightings for each topology in windows along the genome, shown in horizontal order of six scaffolds for nine representative isolates: three soil isolates, three patient isolates, and the three patients that appeared to form a divergent lineage. See Supplementary Figure S14 for the version including all 36 isolates. For both (b) and (c), the bar colors correspond to the topology branch colors in (a).

The *Twisst* results also indicated that both soil and patient-derived *C. immitis* genomes had mosaic architectures. Namely, we found that for a given isolate, no single topology or set of topologies had strong support across the genome (Figure 4A and 4C). Under models specifying three possible topologies, the most common topology for a given isolate accounted for an average of 49.6% (soil) or 49.8% (patient) of genomic windows (Supplementary Table S9). Along these lines, a given dominant topology spanned about 1.9 windows (∼110 kb) for either soil or patient genomes (Figure 4, Supplementary Figure S14 and Table S10). This pattern of mosaic topologies was consistent across bootstrap replicates and an alternative specification of the soil reference groups (Supplementary Results, Supplementary Figures S15–S16, Tables S11– S12). This pattern of genomic mosaicism suggests that different regions of a given soil- or patient-derived *C. immitis* genome have different evolutionary histories, which may be due to ongoing gene flow and/or incomplete lineage sorting.

Across genomic windows, the most common dominant topology estimated by *Twisst* for a given strain tended to match the dominant cluster assignment from ADMIXTURE under *K* = 2 or 3, doing so for 100% of the soil isolates (with this agreement expected to be higher because soil isolates were used to define the reference groups), and 92.9% of the patient isolates (excluding VFI_6, VFI_16, and VFI_27). Interestingly, for the three patient isolates whose genomes exhibited the least sharing with soil references (*i.e.,* were most often positioned on their own branch of the topology rather than as a sister to a soil strain; VFI_6, VFI_16, and VFI_27), ADMIXTURE estimated nearly pure ancestry to the Bakersfield cluster under *K* = 2, but assigned them to their own cluster under *K* = 3–10, with low ancestry to this cluster for all other isolates. This discrepancy highlights the limitations in interpreting a high genome-wide ancestry proportion from ADMIXTURE as close relatedness to that cluster (Supplementary Results).

### No patient isolate could be matched to a point source in the environment

To assess whether any patient *C. immitis* isolates could be plausibly matched to a specific environmental location that we sampled, we calculated pairwise genetic distances between all patient and soil isolates. We found that the minimum genetic distance between any patient and soil isolate was ∼11,300 SNPs (Figure 5)—more than 60-fold greater than the maximum distance between soil isolates from the same burrow (Supplementary Figure S9). This result suggests that the precise sources of patients’ exposure in this study (e.g., the specific rodent burrows) were not included among our sampled soil populations. Yet, given evidence of high genomic divergence and diversity among strains as noted above, the source for a given patient infection could have been anywhere in the region associated with a cluster, either immediately adjacent to one of our collection locations or very far from them.

**Figure 5.**
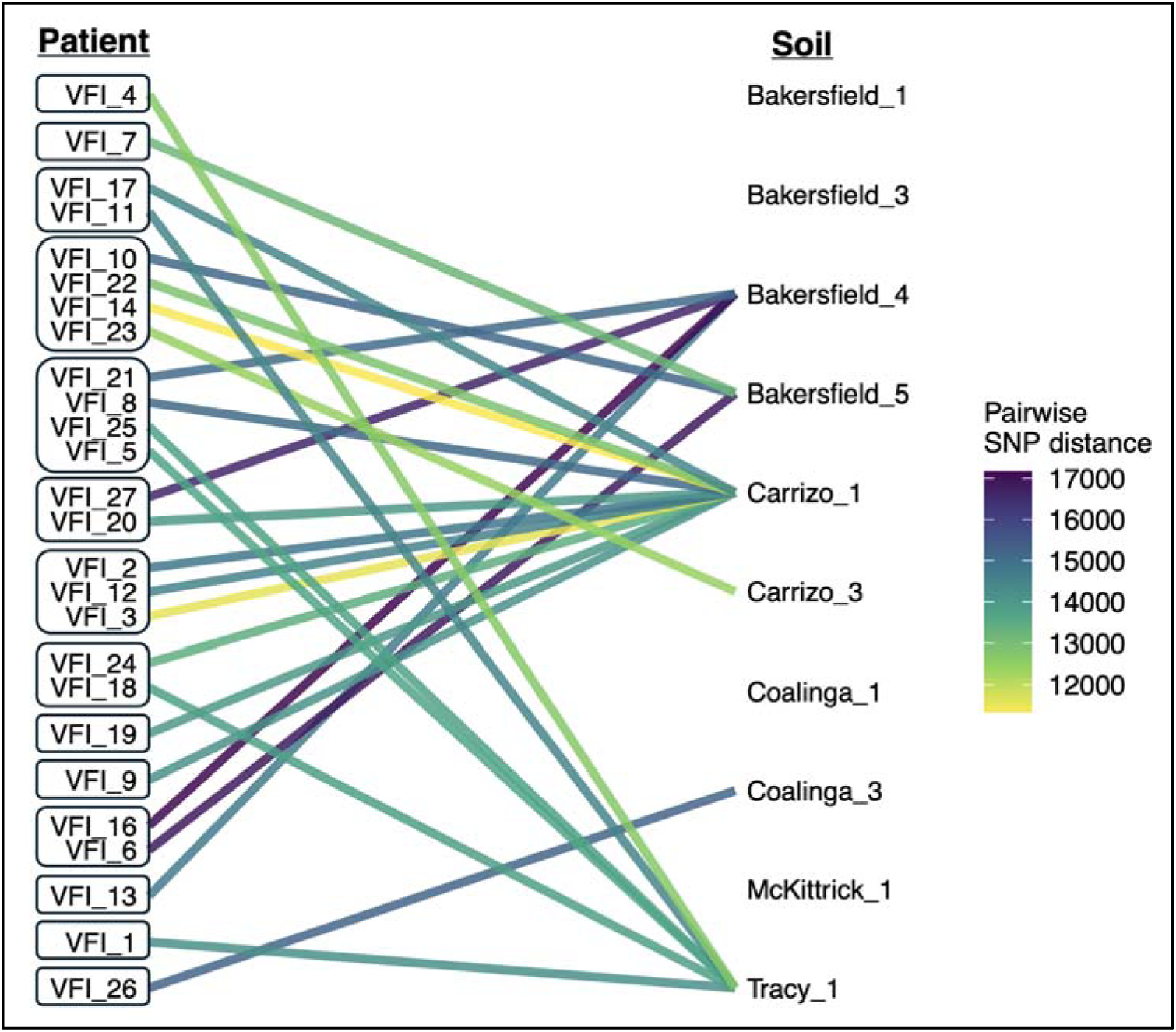
Genetic distance between each patient-derived isolate (left) and the soil-derived isolate (right) to which it had the minimum SNP distance. Patient isolates are grouped based on the zip code of the patient residence, which is not necessarily the geographic source of their exposure. Note that VFI_5 and VFI_25 are *C. immitis* isolates from the same patient; all others are from distinct patients. We include one representative soil isolate per rodent burrow given within-burrow clonality.

Although all patient isolates (n = 26) were derived from individuals with home zip codes in and around the Bakersfield region, the plurality of patient isolates (12 out of 26) were genomically closest to Carrizo soil isolates (Figure 5), despite the Carrizo Plain National Monument site being physically separated from Bakersfield by a low mountain range (the Temblor Range; Figure 1). Further, patient isolates from individuals sharing a home zip code generally did not share a shortest-genomic-distance isolate or population (Figure 5). Thus, our results suggest that a patient’s home zip code constitutes a poor estimate of geolocation for site of exposure.

Two pairs of patient-derived isolates (VFI_5 and VFI_25, VFI_19 and VFI_20) were clonal or near-clonal. These pairs differed by 79 and 29 SNPs, respectively—at or near the level observed for the technical replicates that were known to be identical (47–80 SNPs). VFI_5 and VFI_25 originated from the same patient, collected at the same time from distinct within-host sites. The other clonal clinical pair (VFI_19 and VFI_20) originated from distinct patients residing in distinct Bakersfield zip codes. For both pairs, we retained only one genome per pair when running ADMIXTURE, as noted above for the clonal soil isolates. For VFI_5 and VFI_25, the genetically closest soil match was ∼13,700 SNPs apart and was an isolate from Tracy, approximately 330 kilometers away from the patient’s home zip code in Bakersfield (Figure 5, Supplementary Figure S9). The VFI_19 and VFI_20 clonal pair was genetically closest (∼14,000 SNPs apart) to an isolate from Carrizo, approximately 70–90 kilometers away from the patients’ home zip code and separated by the Temblor Range. These data suggest that, amongst our sampled patient population, exposures to *C. immitis* may have occurred in locations distal from the home zip code; although we could not evaluate source assignments given that the true location of a given patient’s exposure was unknown.

## Discussion

*Coccidioides* is a global priority fungal pathogen and cause of coccidioidomycosis—a serious disease with limited options for control and prevention^36^. As transmission involves inhaling spores aerosolized from the soil, understanding how fungal populations are distributed in the environment and where people are becoming exposed is critical for minimizing infection risk. In this study, we generated the first whole genomes of *C. immitis* from the endemic environment— isolated from soils drawn from rodent burrows across California’s San Joaquin Valley—and analyzed these alongside novel patient-derived genomes from a similar region and time period (Figure 1). We found that environmental *C. immitis* is structured at regional scales, with extensive allele sharing between clusters, and high genomic diversity partitioned into effectively clonal lineages in individual rodent burrows. Further, the majority of patient-derived isolates shared full ancestry with a single cluster containing soil isolates, although no patient exposure could be matched to a specific soil source sampled in our study.

Our study provides evidence of population structure among environmental strains of *C. immitis* within the endemic region. Prior research based on patient-derived genomes has indicated separation between populations from the San Joaquin Valley, and San Diego and Mexico^37,38^, yet finer-scale investigation was not possible as the source location of a patient’s exposure is typically unknown. In this study, we generated *C. immitis* genomes from precisely georeferenced rodent burrows, which we targeted for sampling given prior evidence that *Coccidioides* is closely associated with small burrowing mammals^3,4,18,22^. This enabled identification of multiple genomic clusters within the San Joaquin Valley, with primary separation of environmental strains from Bakersfield versus other collection locations (i.e., Carrizo, Coalinga, McKittrick, and Tracy), and further separation within the non-Bakersfield group in models assuming additional clusters (Figure 2). The clusters were moderately differentiated (F_ST_ of 0.15–0.23) rather than genetically isolated, as soil isolates were broadly separated by collection location on the phylogenetic tree but did not form reciprocally monophyletic groups, and window-based analyses indicated that each of our isolates shared ancestry across clusters (Figure 4). Taken together, these findings are consistent with ongoing gene flow between the inferred clusters (or ancestral variation from a recent divergence, which we cannot rule out).

Within this regionally structured environmental pool, we found that *C. immitis* diversity was partitioned into clonal or nearly clonal lineages within individual rodent burrows. Specifically, isolates from the same burrow differed by fewer than 200 SNPs (on par with our sequencing controls), and 52% of all SNPs segregating among the soil samples were private to a single burrow, indicating that each burrow lineage harbors substantial unique variation. Yet whereas isolates from a given burrow were clonal or near-clonal, those from adjacent burrows—for example, separated by less than 200 meters—differed by a minimum of over 13,000 SNPs (Figure 3). Notably, this genomic distance is as much as that observed for some pairs separated by tens to hundreds of kilometers, including pairs on opposite sides of a low mountain range (the Temblor Range). These findings are consistent with a scenario in which *Coccidioides* spores can be dispersed (e.g., via wind, mechanical disturbance, or natural disasters^39,40^)—including up to hundreds of kilometers away in some instances^20,41^—and then establish clonal or nearly clonal lineages at deposited locations, if conditions are favorable. Consequently, for a given population, as determined by inference schemes of the kind we use here, its geographic extent would depend on past dispersal events. Dispersal over long distances, followed by lineage maintenance, has previously been postulated for instances of *Coccidioides* range expansion, as clonal lineages have been identified in Washington State^28^ and Venezuela^42^—areas outside of the historically endemic range.

Our results indicated that patient-derived isolates—generated from individuals residing in or near Bakersfield, California—were indistinguishable from a random draw from the environmental pathogen pool. In particular, we detected no overall differentiation between soil- and patient-derived genomes (F_ST_ of 0.013, permutation p = 0.14), suggesting a lack of clear separation between *Coccidioides* populations in the environment and those inhaled by a human host. Further, across genomic windows, patient-derived isolates were consistently estimated to share ancestry with sampled soil populations rather than form their own branches (Figure 4). The one exception was a set of three patient-derived isolates that shared two identical, likely derived deletions (containing six genes, each of unknown function) and formed a divergent lineage (Supplementary Figure S10). We further found that our patient- and soil-derived genomes harbored similar levels of genomic diversity (Supplementary Table S7), consistent with the absence of a strong genetic bottleneck to human infection. However, we note this finding is not conclusive given the unknown exposure source location for our patient-derived isolates and the relatively small sample sizes.

Despite overall genomic similarity at the population level, none of the patient-derived *C. immitis* strains could be matched to a point source in the environment. In particular, the nearest match between patient and soil isolates was ∼11,300 SNPs, suggesting that we did not collect samples from the precise locations at which patients in this study were exposed. This result may be expected given our findings that environmental populations vary substantially at sub-kilometer scales (i.e., by tens of thousands of SNPs), and that we estimated having captured at most 57% of the genomic variation segregating among our soil populations. Thus, we cannot rule out that the source of patient exposures was nearby to our sampled soils. Notably, our Bakersfield soil samples were collected along the Kern River Parkway Bike Trail, in close proximity to a dense urban area (Figure 1). Yet, no soil-derived isolate from the Bakersfield collection site was within 13,000 SNPs of a patient-derived isolate. By contrast, a prior study linked a case of coccidioidomycosis to exposure at a specific park in eastern Washington state, as the patient-derived isolate was within 3 SNPs of soil-derived isolates collected from that location^28^ (though we note that SNP counts are not directly comparable between studies due to variation in sequencing depth and bioinformatic filtering thresholds^43,44^). The environmental surveillance and genomic linkage conducted in that study were critical for establishing exposure in Washington, leading to an updated understanding of the endemic distribution for *C. immitis,* which was previously considered limited to California.

Our study provides the first insight into the structure and diversity of environmental *C. immitis* populations. Our detection of population structure among *C. immitis* within the San Joaquin Valley suggests that strategies to identify general regions of human exposures to pathogenic spores within this highly endemic region may be possible. Yet the high diversity and gene flow in *Coccidioides* we identified also highlight the need for increased collection and sequencing of environmental strains of *Coccidioides* beyond that conducted in this study. Such efforts are becoming more tractable, as approaches for targeted DNA capture and enrichment from mixed samples, which avoid the need for high-containment facilities, have recently been developed^45^. Further, it remains unknown whether all environmental sources of *Coccidioides* pose an equal human health risk, or whether certain populations harbor heightened capacity for virulence. We could not directly investigate this question in the present study as the origin of exposure was unknown for our patient-derived isolates; thus, any genomic comparisons could have been readily confounded by demography, particularly given the spatially stratified nature of our soil samples. Future work using experimental approaches, including conducting comparative phenotypic assays on soil- and patient-derived isolates, and/or tracking isolate diversity throughout a time course of infection, would be valuable for identifying underlying mechanisms of virulence. Given the rapid rise of coccidioidomycosis cases and the limited options for prevention and treatment, a better understanding of human exposure pathways is critically needed.

## Materials and Methods

### Sample collection and isolation

We isolated *Coccidioides* from soil samples collected across California between 2018–2025 for this study or as part of prior studies by our group^3,32–34^. These soil samples had previously tested positive for the presence of *Coccidioides* DNA using the CocciEnv qPCR assay^46^ (Supplementary Methods). From these qPCR-positive soil samples, we used the “double pour” method for *Coccidioides* culturing^31,47^. In brief, each soil sample was first mixed with water and allowed to settle, then 0.5 mL of the topmost supernatant was plated, followed by two additions of yeast extract agar fortified with cycloheximide, chloramphenicol, and streptomycin (Supplementary Methods). Cultures were grown for one week at 30°C under BSL3 containment, then colonies with potential *Coccidioides* morphology were tested via PCR and Sanger sequencing of the ITS2 region. All putative *Coccidioides* colonies were then transferred to a fresh agar plate, and subcultures were grown for an additional week, then re-tested via ITS2 sequencing (Supplementary Methods).

Patient isolates of *Coccidioides* were obtained from the Valley Fever Institute at Kern Medical in Bakersfield, California. These isolates were derived from clinical specimens collected from patients between 2024–2025 as part of routine diagnostic and clinical evaluation. Specimen types collected included sputum (n = 6), bronchial washings (n = 3), tissue (n = 6), abscess (n = 5), fluid (n = 3), and cerebrospinal fluid (CSF; n = 3). Patient specimens were inoculated onto individual Sabouraud dextrose agar slants and incubated at 30°C for a total of four weeks.

### DNA extraction and whole-genome sequencing

After confirming isolate identity, we extracted DNA from each isolate using the ZR Quick-DNA™ Fungal/Bacterial Miniprep Kit according to manufacturer’s instructions (Zymo Research Corp., Irvine, CA, USA). For each sample, we submitted 100-500 ng gDNA to the California Institute for Quantitative Biosciences (QB3) at UC Berkeley for QC and standard library preparation using the Illumina DNA prep kit. Libraries were then pooled and sequenced as 150-bp paired-end reads on a NovaSeq 6000 using an S4 flow cell.

### Legacy *Coccidioides* genomes

In addition to the novel soil and patient genomes generated in this study, we included previously published patient-derived *C. immitis* genomes (hereafter “legacy patient genomes”). Metadata for these isolates are less specific than for our novel isolates, but the reported origins spanned western North America and Mexico with collection dates between the 1970s and 2019 (metadata in Supplementary Table S13).

### Read trimming and variant calling

All sequence data, including those from new and previously published genomes, were subject to consistent downstream bioinformatic processing^44^. Namely, raw sequence reads were first trimmed for low-quality bases and Illumina adapters using *Trimmomatic v0.39*^48^ (Supplementary Methods). Read lengths were then normalized to 75 bases, to account for variation in sequence read length across previously published genomes, which can introduce biases in downstream steps^49,50^ (Supplementary Methods). High-quality reads were then aligned to the *C. immitis* reference genome (*C. immitis* RS GCF_000149335.2) using BWA-MEM2 (version 2.2.1)^51^. Duplicate reads were marked and removed using picard (version 3.0.0)^52^, and variants were called on individual samples using HaplotypeCaller, followed by joint genotyping on all genomes (n = 86) using GenotypeGVCFs from GATK (version 4.5.0)^53^ (Supplementary Methods). This all-sample VCF file was used in phylogenetic analyses; a subset containing just our novel genomes was used in all other analyses. Variants were flagged and removed using VariantFiltration and SelectVariants from GATK if they were either multi-allelic or met any of the following criteria: Depth of Coverage < 10, Quality by Depth < 2.0, Fisher Strand Bias > 60.0, Mapping Quality < 40.0, Minor Allele Frequency < 0.05, or Phred-scaled Quality Score < 30. Further, to avoid potential confounding by genotype missingness in downstream analyses, only samples for which 90% of the genome was covered at >10x depth were retained. Lastly, to assess the extent of residual technical variation, we repeated the full workflow from sub-culturing through variant calling for three of the soil-derived samples. These technical replicates enabled us to estimate an approximate threshold of pairwise SNP differences expected among clonal isolates.

### Recombination

Given evidence of potential non-independence among SNPs in the genotype matrix (Supplementary Figure S17), we investigated the extent of recombination among our soil and patient isolates. We calculated both the standardized index of association (*r̅_d_*, which measures the extent to which alleles along the genome are independent or linked^54^, and the linkage disequilibrium decay distance (Supplementary Methods). To obtain an LD-corrected SNP set for use in certain downstream analyses, we used a SNP pruning algorithm implemented in plink (version 1.9)^55^. Herein, SNPs were iteratively pruned such that no two SNPs had an r^2^ > 0.5 within a 50-SNP window, advanced by 5 SNPs at a time. A total of 12,834 (all 86 samples) or 11,718 SNPs (our soil and patient samples only) were retained after this step.

### Genomic diversity

We assessed genomic diversity within soil and patient isolates by calculating the following metrics: the number of segregating sites (S), Watterson’s estimator (θ_w_), and mean nucleotide diversity (π) (Supplementary Methods). In these calculations, we excluded one isolate from any pairs that appeared nearly clonal (i.e., <200 SNP differences) to avoid downward bias in diversity estimates. We also investigated genomic diversity among the soil-derived isolates by examining the SNP accumulation curve across burrows and calculating the Chao2 estimator^56^—a lower bound estimate of the total number of SNPs segregating in our soil populations (Supplementary Methods).

To explore the potential drivers and bottlenecks of *C. immitis* genetic diversity, we investigated the distribution of mating type loci among all samples. Namely, *Coccidioides* is known to have a single mating type locus with one of two forms (‘idiomorphs’)—*MAT1-1* and *MAT1-2*—and sexual reproduction can only occur between isolates with different idiomorphs^57^. We determined the mating type of each isolate using idiomorph-specific marker sequences for *Coccidioides* using the alpha-box gene (EF472259.1) for *MAT1-1* and the HMG domain (EF472258.1) for *MAT1-2*^38,42^.

### Population structure

We investigated population structure among our isolates using two complementary approaches. First, we conducted a principal components analysis (PCA) on the filtered, bi-allelic variant matrix using *pcadapt* in R^58^. We examined the proportion of variance explained by each axis and visualized the clustering of samples, including by source (soil versus patient), year of collection, and patient residence within or outside of Bakersfield zip codes. Next we estimated ancestry proportions for each isolate using ADMIXTURE (version 1.4.0)—a maximum likelihood approach for inferring clusters of genomic ancestry given *K* ancestral populations^59^—on the LD-pruned SNP set. We estimated ancestry across *K* = 2 to 10, using 20 replicate runs per *K*, and examined cross-validation (CV) errors across *K* and across replicates. We then calculated mean genome-wide weighted F_ST_ (Weir and Cockerham) between inferred soil populations.

### Phylogenetic analysis

We investigated the evolutionary relationships among our novel *C. immitis* isolates, as well as the legacy patient genomes by inferring a maximum-likelihood phylogenetic tree from our multi-sample VCF using IQ-TREE (version 3.0.0)^60^. We used the best-fit nucleotide substitution model determined by ModelFinder^61^ with ascertainment bias correction, and assessed branch support using 1,000 ultrafast bootstrap replicates^62^ and 1,000 Shimodaira-Hasegawa approximate likelihood ratio test (SH-aLRT) replicates^63^. For the rooted version of the tree, we used *C. posadasii* (Silveira strain, GCA_018416015.2) as the outgroup.

To investigate how estimated phylogenetic relatedness varied along the genome, we conducted a sliding-window topology analysis using *Twisst* (topology weighting by iterative sampling of subtrees)^35^, focusing on our novel soil and patient isolates. Herein, a full tree containing all samples is first estimated for windows along the genome. From this full tree, smaller sub-trees consisting of one isolate per group are repeatedly sampled. The groups consist of the focal isolate, user-defined reference groups, and an outgroup. The possible topologies have the focal isolate sister to a reference group, or on its own branch closer to the outgroup. The support for a given topology at that window is defined as the proportion of sub-trees reflecting that topology.

We used *C. posadasii* as the outgroup, and the soil *C. immitis* isolates as reference groups, using one representative isolate per burrow given the within-burrow clonality. In the main model specification, we defined two soil reference groups based on the inferred genomic clusters from ADMIXTURE under both *K* = 2 and 3, which split soils from Bakersfield and all other sites. This allowed for three possible topologies (i.e., sister to Bakersfield, sister to non-Bakersfield, or own-branch). In an alternative model specification, we defined three soil reference groups using clusters inferred under ADMIXTURE at *K* = 4, which split the non-Bakersfield group into the southwestern Central Valley (Carrizo and McKittrick collection sites), and the western Central Valley (Coalinga and Tracy collection sites). This allowed for 15 possible topologies (Supplementary Methods). We considered each soil or patient isolate, in turn, as focal, leaving out the soil isolate from the reference group when it was focal.

Trees were estimated using maximum likelihood in RAxML (version 8.2.12)^64^ under the GTRCAT model with a window size of 100 SNPs. As a sensitivity analysis, we repeated the analysis under alternative specifications of the soil reference groups (Supplementary Methods). To assess uncertainty in the per-window trees, we conducted 100 bootstrap iterations wherein the 100 SNPs defining a given window were randomly sampled with replacement, followed by per-window tree estimation and topology weighting. Reported values and ranges, where supplied, are the mean and 95% confidence intervals from the bootstrap distribution.

## Ethics statement

Isolation and extraction of *Coccidioides* was conducted in Biosafety Level 3 (BSL3) facilities in accordance with Environmental Health and Safety guidelines under Biological Use Authorization #223. This study received ethical approval from the Kern Medical Institutional Review Board (IRB # 25055).

## Data availability

*Coccidioides immitis* sequence data generated in this study are available under NCBI BioProject PRJNA1522484. Previously published sequence data are accessible using the SRA Accession numbers listed for each in Supplementary Table S2. All other metadata are reported in the Supplement.

## Code availability

All scripts used for bioinformatics or downstream analyses are available at: https://github.com/lcouper/CocciGenomics/.

## Acknowledgements

We thank Matthew Morales, Ashley Itogawa, and Dr. Bridget Barker for providing training in *Coccidioides* culturing techniques, which was invaluable to this study. This work was supported by the National Institutes of Health under grant number R01AI148336 awarded to JVR, and U19AI166798 and R21AI193607 to RBB. LIC was supported by a National Science Foundation Postdoctoral Research Fellowship in Biology.

