## Supplementary Information for "Population structure of human-infecting and environmental isolates of *Coccidioides immitis*, the fungus causing Valley fever"

**Supplementary Methods**

1. Environmental sample collection and testing
2. *Coccidioides* isolation
3. Read trimming and variant calling
4. Recombination
5. Genomic diversity
6. Topology weighting

**Supplementary Results**

1. Technical replicates
2. Isolation by distance
3. Population structure under alternative specification
4. Topology weighting under different reference group specifications

**Supplementary Figures S1–S17**

**Fig S1.** *Coccidioides* culture success by year and month of collection

**Fig S2.** CV error and percent explained across *K* values

**Fig S3.** ADMIXTURE results across *K* = 5–10

**Fig S4.** ADMIXTURE results under alternative specification

**Fig S5.** K-means clustering on PCA

**Fig S6.** PCA by year of collection and patient residence

**Fig S7**. Maximum likelihood phylogenetic tree

**Fig S8.** LD-decay curves

**Fig S9.** Heatmap of pairwise SNP distances among all soil and patient isolates

**Fig S10**. Genotype matrix surrounding focal deletion

**Fig S11.** Location of SNPs in technical replicates and same-burrow samples

**Fig S12.** SNP accumulation curve across burrows

**Fig S13.** Genetic vs geographic distance between isolates or sampling sites

**Fig S14.** Window-based topology weights under three possible topologies

**Fig S15**. Average topology weights under 15 possible topologies

**Fig S16.** Window-based topology weights under 15 possible topologies

**Fig S17.** Scaffold-level genotype matrix heatmap

**Supplementary Tables S1–S13**

**Table S1.** Source metadata for *C. immitis* genomes

**Table S2.** Sequence metadata

**Table S3.** F_ST_ values for soil populations under varying *K*

**Table S4.** $\bar{r}_{d}$ and LD half-decay distances

**Table S5.** Number of private alleles per burrow

**Table S6.** Geographic and genomic distances across scales

**Table S7**. Diversity metrics

**Table S8**. Mating type loci assignments

**Table S9.** Dominant topology weights under three possible topologies

**Table S10.** Dominant topology switch rates under three possible topologies

**Table S11.** Dominant topology weights under 15 possible topologies

**Table S12.** Dominant topology switch rates under 15 possible topologies

**Table S13.** Source metadata for legacy *C. immitis* genomes

**Supplementary References**

**Supplementary Methods**

Environmental sample collection and testing

Soil samples were collected from the San Joaquin Valley of California between 2018–2025 for the purposes of this study or prior work by our group^1–4^. Samples were predominantly taken from rodent burrows and adjacent surface soils. All samples that resulted in successful isolation and inclusion in this study came from rodent burrows. DNA was extracted using the Qiagen PowerSoil kit (QIAGEN, Valencia, CA, USA) according to the manufacturer’s instructions. Samples were then tested for the presence of *Coccidioides* DNA using the CocciEnv qPCR assay, using four replicates per sample. Samples in which at least three out of four wells had a Ct value < 40 were deemed positive, as in our prior work^1,2,4^.

*Coccidioides* isolation

Using the positive soil samples, we attempted culture-based isolation of *Coccidioides* following the “double-pour” method^5,6^. Herein, 2.5 g of soil was combined with 20 mL sterile DI water, thoroughly vortexed, then allowed to settle at room temperature. After one hour, 0.5 mL of liquid from the surface above the settled soil was added directly to the center of a Petri dish. Immediately thereafter, 15 mL of growth media was added to the Petri dish. Growth media consisted of 3:4:200 g Difco^TM^ yeast extract, Difco^TM^ agar, and DI water that was autoclaved, and then fortified with 50 ppm of chloramphenicol, 0.5 mg/mL cycloheximide and 0.32 mg/mL streptomycin sulfate.

After 2-3 minutes, during which time the media had partially hardened in the plate, a second round of 15 mL growth media was added. Once the second round of media had solidified, Petri dishes were transported in secondary containment to the BSL3 facility at UC Berkeley where they were maintained at 30°C in an incubator. After seven days of growth, any colonies with *Coccidioides*-like morphology (fluffy, white) were harvested and identified by Sanger sequencing of the ITS2 region.

Any colonies identified as *Coccidioides* were then sub-cultured on a fresh YEA plate with antibiotics. After one week of sub-culture at 30°C, colonies were then re-tested via Sanger sequencing. After confirmation of *Coccidioides i*dentity, DNA was extracted from subcultures using the ZR Quick-DNA™ Fungal/Bacterial Miniprep Kit according to manufacturer’s instructions (Zymo Research Corp., Irvine, CA, USA). The resulting DNA concentrations were measured via Qubit, and 100-500 ng of genomic DNA was submitted to the California Institute for Quantitative Biosciences (QB3) at UC Berkeley for quality control, library preparation, and DNA sequencing on an Illumina NovaSeq 6000 with an S4 flow cell.

Read trimming and variant calling

We trimmed raw sequence reads for low-quality bases and Illumina adapters using *Trimmomatic v0.39*^7^ with the following parameters: ILLUMINACLIP:TruSeq3-PE.fa:2:30:10 LEADING:3 TRAILING:3 MINLEN:35 SLIDINGWINDOW:4:15. After trimming low quality base calls and adapters, we normalized the read lengths for all samples to 75—the lowest sequence read length used across the legacy patient genomes considered in this analysis. That is, while all of our environmental samples were sequenced as 150-bp paired-end reads, prior genomes were sequenced at 75-300-bp paired-end reads. As variation in read length can impact alignment, mapping quality, and variant-calling, introducing biases in downstream steps^8,9^, we sought to minimize this effect through read length normalization. We note that all genomes generated as part of this study were sequenced on the same Illumina platform (NovaSeq 6000), thus reducing any additional biases introduced by variation in sequencing technology.

High-quality reads were then aligned to the *C. immitis* reference genome (*C. immitis* RS GCF_000149335.2) using BWA-MEM2 (version 2.2.1)^10^, following masking repetitive regions in this reference genome using RepeatMasker (version 4.1.0)^11^ and the *C. immitis* repeat elementary library^12^. Read groups were then added to each sample using *picard* (version 3.0.0)^13^ to enable downstream analyses to account for variation in sequencing platform and library preparation. Duplicate reads were marked and removed using *picard*, and variants were called on individual samples using HaplotypeCaller in GVCF mode with ploidy set to 1, followed by joint genotyping on all genomes (including the legacy genomes; n = 86 in total) using GenotypeGVCFs from *GATK* (version 4.5.0)^14^. Variants were then filtered as described in the main text.

Recombination

We investigated the extent of recombination among our isolates by calculating the standardized index of association ($\bar{r}_{d})$^15^, which captures the extent to which alleles at different sites along the genome are independent. We used poppr^16^ to estimate $\bar{r}_{d}$ among groups of isolates (soil only, patient only, or all), using 200 random draws of 100 SNPs. To avoid confounding based on sample size differences (i.e., 20 patient samples versus 10 soil samples after removing near-clones), we repeated $\bar{r}_{d}$ estimation on 50 random subsamples from the patient isolates to match the soil sample size. We also excluded any potentially clonal isolate pairs (i.e., those with pairwise SNP distance <200 SNPs) from this estimation to avoid falsely inflating $\bar{r}_{d}$ due to clonality.

We also estimated r^2^ between pairs of SNPs as a function of genomic distance (i.e., linkage disequilibrium decay). Namely, we estimated r^2^ in 100 kb windows and identified the distance at which r^2^ was half of its initial value. As with $\bar{r}_{d}$, we estimated r^2^ among groups of isolates (soil only, patient only, or all), while removing any potentially clonal isolates.

Genomic diversity

We assessed genomic diversity among our samples by calculating several genome-wide diversity metrics including: the number of segregating sites (S), Watterson’s estimator (θ_w_), and mean nucleotide diversity (π). We calculated these metrics for the following sets of samples: 1) all soil samples, 2) all patient samples, 3) the soil and patient samples, and 4) all samples including the ‘legacy’ genomes. These metrics were calculated from the shared filtered VCF and normalized by the number of callable sites. We note that the shared VCF was filtered to remove variants with a minor allele frequency < 0.05, thus causing downward bias in the reported diversity metrics. Nucleotide diversity was calculated using *vcftools* (version 0.1.16)^17^. Technical replicates and near-clones (i.e., differing by <200 SNPs from another isolate) were excluded from these calculations.

For the soil isolates, we also calculated pairwise SNP differences and examined the evidence for isolation by distance by conducting a Mantel test using the *vegan*^18^ package in R. Specifically, we calculated the correlation between genomic distance (here, the number of SNP sites that differ) and the natural log of geographic distance (great-circle distance) between pairs of soil isolates, and determined statistical significance using 9,999 permutations of the genomic distance matrix. We conducted this analysis using soil isolates after removing near-clones (n = 10). We also estimated our power to detect IBD via simulation, in which we held the observed geographic distance matrix fixed but generated alternative genomic distances with a specified correlation to geographic distance. We repeated this across a range of correlations and evaluated each using the same permutation test applied to the observed data.

We further investigated genomic diversity among the soil-derived isolates by examining the SNP accumulation curve across the ten burrows and calculating the Chao2 estimator^19^—a lower bound estimate of the total number of SNPs segregating in our soil populations. In our context, the Chao2 estimator is calculated based on the presence or absence of specific SNPs across multiple burrows, relying primarily on SNPs unique to a single burrow ($Q_{1})$ or found in only two burrows ($Q_{2})$. Specifically, it is expressed as:

$S_{Chao2}= S_{obs}+ \frac{m-1}{m} \times\frac{Q_{1}^{2}}{2 Q_{2}}$

where $S_{obs}$ is the observed number of segregating SNPs across all burrows, and *m* is the total number of burrows sampled (here, 10).

Topology weighting

We investigated phylogenetic relationships among our soil and patient isolates using topology weighting by iterative sampling of subtrees (*Twisst*)^20^, which estimates the support for every possible topology along sliding windows of the genome. We used our soil isolates as reference groups, using one representative isolate per burrow given the within-burrow clonality. We defined either two (main text) or three (alternative specification) soil reference groups, using the genomic clusters inferred by ADMIXTURE under *K* = 2 and 3 (two groups) or *K* = 4 (three groups). We used *C. posadasii* (Silveira strain) as the outgroup. Specifically, under two soil reference groups, we defined groups as ‘Bakersfield’ (which included 4 soil isolates from Bakersfield) and ‘non-Bakersfield’ (which included 6 soil isolates from Coalinga, Tracy, Carrizo, and McKittrick). Under three soil reference groups, we defined groups as ‘Bakersfield’, ‘southwestern Central Valley’ (2 Carrizo, 1 McKittrick), and ‘western Central Valley’ (2 Coalinga, 1 Tracy). Each soil or patient isolate was then, in turn, considered the ‘focal’ isolate. If a soil isolate was focal, it was not included in a reference group.

Given two or three soil reference groups, there were three or 15 possible topologies, respectively (main text Figure 4). The weight for a given topology is calculated as the proportion of sub-trees that match it, with sub-trees formed by taking one individual per group from the full tree. Trees were estimated using maximum likelihood in *RAxML* (version 8.2.12)^21^ under the GTRCAT model with a window size of 100 SNPs. This resulted in 531 windows. We assessed uncertainty in the per-window trees by generating 100 bootstrap replicates wherein the 100 SNPs defining a given window were randomly sampled with replacement, followed by per-window tree estimation and topology weighting.

We investigated how group membership assignments varied between ADMIXTURE and *Twisst*. We note that these two methods are based on different inputs (LD-pruned genome-wide SNPs versus unpruned SNPs partitioned into windows) and inference goals (i.e., estimating individual ancestry proportions from allele frequencies versus estimating relative weight for varying topologies), thus the expectation was not for perfect agreement. Rather, the goal here was to assess agreement in interpretation between distinct approaches. However, the *Twisst* reference groups were based on the clusters inferred from ADMIXTURE, thus the two methods are not independent, and agreement for the soil isolates is expected to be high. The assignments for the clinical isolates are thus the comparison of interest.

**Supplementary Results**

Technical replicates

To identify the extent to which sequence variation reflected random error rather than true biological signal, we generated a technical replicate for each of three soil-derived genomes (specifically Bakersfield_1, Bakersfield_3, and McKittrick_1). For each, we started a new sub-culture of the original isolate. We then extracted genomic DNA and submitted samples for QC, library preparation, and sequencing at California Institute for Quantitative Biosciences (QB3) at UC Berkeley. Technical replicates underwent the same bioinformatic steps as all other samples. In doing so, we found that the technical replicates differed from their matched isolate by 47, 53, or 80 SNPs. These SNPs were more common in certain positions of the genome, particularly within a few kilobases of repeat-masked sequences, suggesting they may be due to challenges in mapping to the reference (Supplementary Figures S11 and S17).

Isolation by distance

We tested the relationship between genomic similarity and physical separation of collection location using a single member from each burrow and found no significant isolation by distance (IBD) relationship (Mantel r = 0.221, p = 0.095; Supplementary Figure S13). However, we noted that our power to detect IBD was limited by the number of independent burrow lineages (i.e., 45 pairwise comparisons given 10 burrow lineages). Specifically, from our power calculations, we estimated that we had 18% power to distinguish the observed IBD effect size from zero.

Population structure under alternative specification

As described in the main text, we investigated population structure using principal components analysis (PCA) and ADMIXTURE^22^ across *K* = 2–10. Given our finding that several isolates had a clonal mate (including all isolates from a given rodent burrow, and two pairs of clinical isolates), we included only one genome from a given clonal set when running ADMIXTURE to avoid biasing estimation towards stronger population structure. Results under this specification are reported in the main text. As an alternative specification, we included all isolates (i.e., retaining all those from a clonal set) and ran ADMIXTURE under identical parameters. As expected, we found that doing so led to stronger population structure for the soil isolates in particular, with more soil isolates estimated to have pure ancestry to a given cluster (Supplementary Figure S4).

Topology weighting under different reference group specifications

We conducted the *Twisst* (topology weighting by iterative sampling of subtrees) analysis using either two or three soil reference groups. Results generated using two soil reference groups (‘Bakersfield’ and ‘non-Bakersfield’) are described in the main text, and results generated using three soil reference groups (‘Bakersfield’, ‘southwestern Central Valley’, and ‘western Central Valley’) are as follows. A given soil-derived isolate was placed in a clade with a soil reference population in 82.7% of its genomic windows on average (Supplementary Table S11). A given patient-derived isolate was placed with a soil reference population across 82.2% of its genomic windows on average. As under the main specification, this result excludes three patient isolates (VFI_6, VFI_16, and VFI_27), each of which was more commonly placed on its own branch. Specifically, the three isolates were placed on their own branch across 42.0% of windows on average.

The *Twisst* results indicated that both soil and patient-derived *C. immit*is genomes had a mosaic architecture, as under the main reference group specification. Namely, the most common dominant topology (out of 15) for a given isolate accounted for an average of 17.6% (soil) or 15.6% (patient) of the genomic windows (Supplementary Figure S15, Table S11). Further, the most common dominant topology spanned about 1.2 windows (~69.8 kb) for either the soil or patient isolates (Supplementary Figure S16, Table S12).

The dominant topology estimated by *Twisst* matched the dominant cluster assignment from ADMIXTURE under *K* = 4 for 100% of the soil isolates and 59.2% of the patient isolates. We note the soil agreement is expected to be high as soil isolates were used to define the reference groups (although a given soil isolate was removed from the reference groups when it was focal). When the assignments disagreed, *Twisst* tended to more frequently place isolates in a clade with the southwestern Central Valley group (containing the Carrizo and McKittrick isolates) while ADMIXTURE assigned them to the western Central Valley cluster (containing the Coalinga and Tracy isolates). This accounted for 56.9% of disagreements.

**Supplementary Figures**

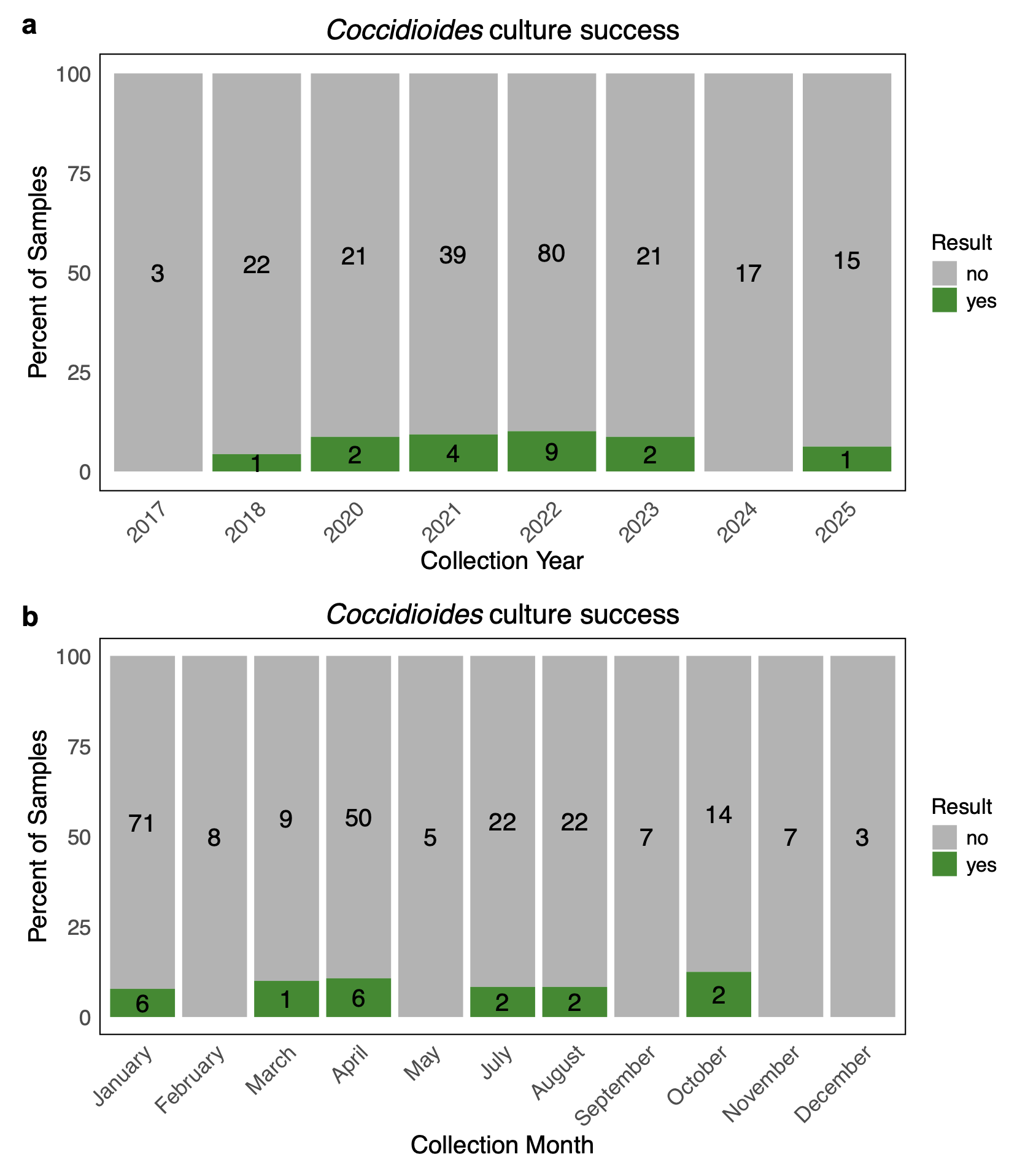

**Supplementary Figure S1.** *Coccidioides* culture success by the **(a)** year or **(b)** month of soil sample collection. Culturing was only attempted on soil samples with evidence of *Coccidioides* DNA based on positive qPCR results. Note that not all *Coccidioides* isolates were successfully sequenced.

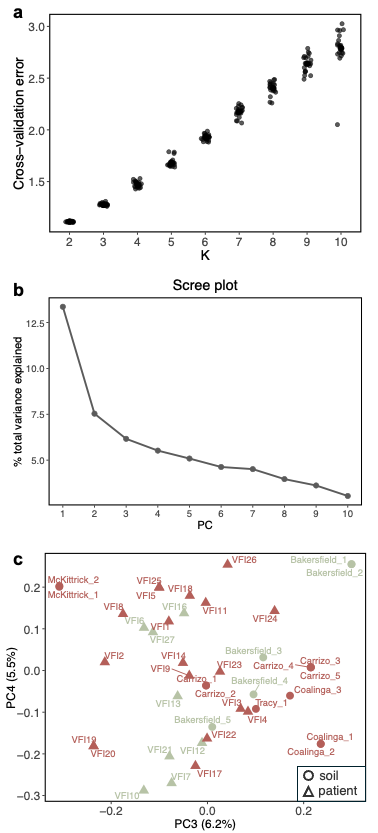

**Supplementary Figure S2.** **(a)** Cross-validation error across *K* = 2–10. Each point denotes the CV error from a single run (n = 20 per *K*). **(b)** Scree plot showing the percent of variance explained by each principal component in the PCA. **(c)** PCA of genomic variation using the third and fourth principal components. Soil-derived isolates (circles) and patient-derived isolates (triangles) are colored by their dominant cluster assignment under *K* = 2 and 3. The percent variance explained by each axis is shown in parentheses.

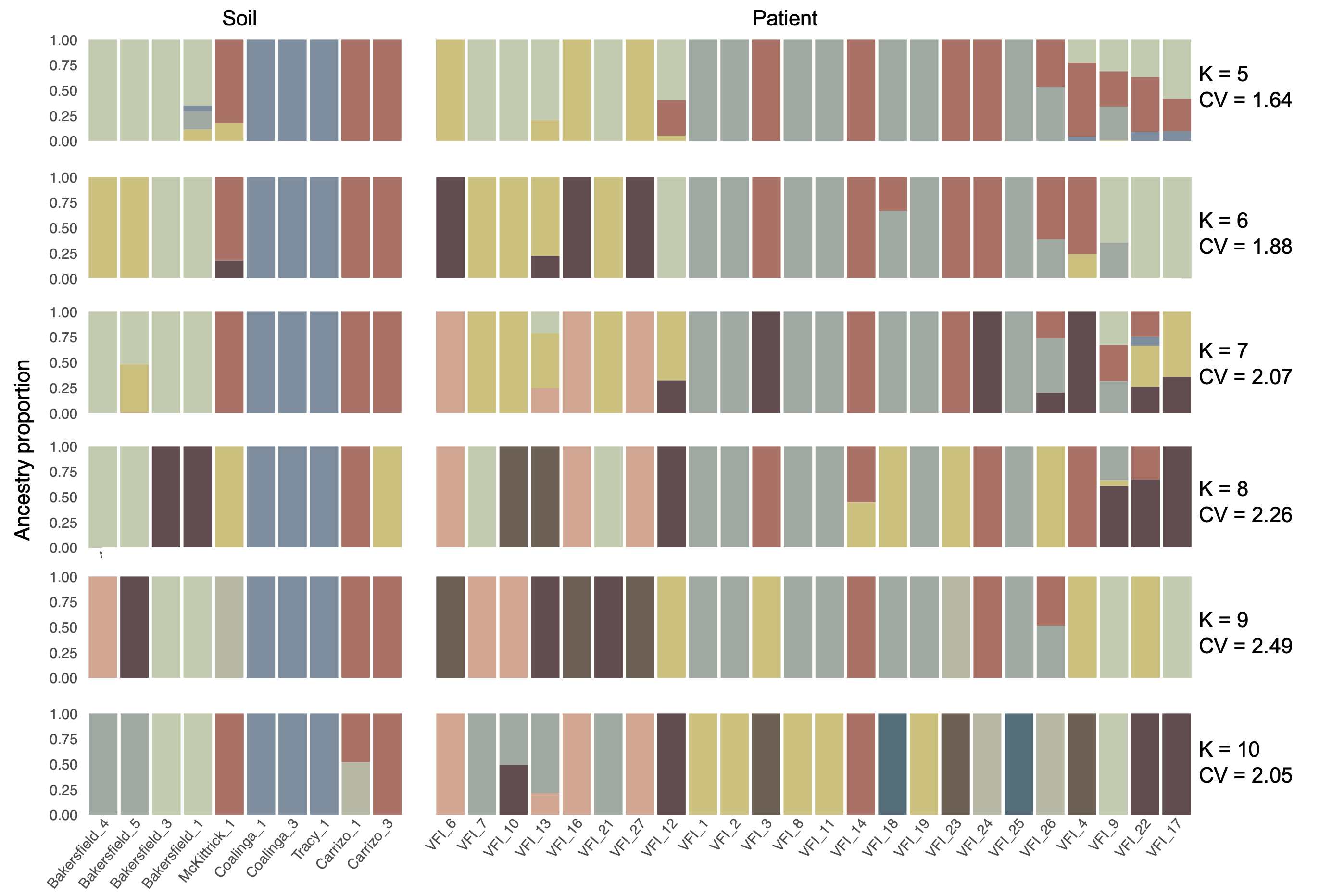

**Supplementary Figure S3.** Stacked bar plots showing ancestry proportions estimated by ADMIXTURE under *K* = 5–10. Each bar represents an isolate, with soil-derived on the left and patient-derived on the right. The mean cross-validation error for a given *K* is listed on the right side of the plot.

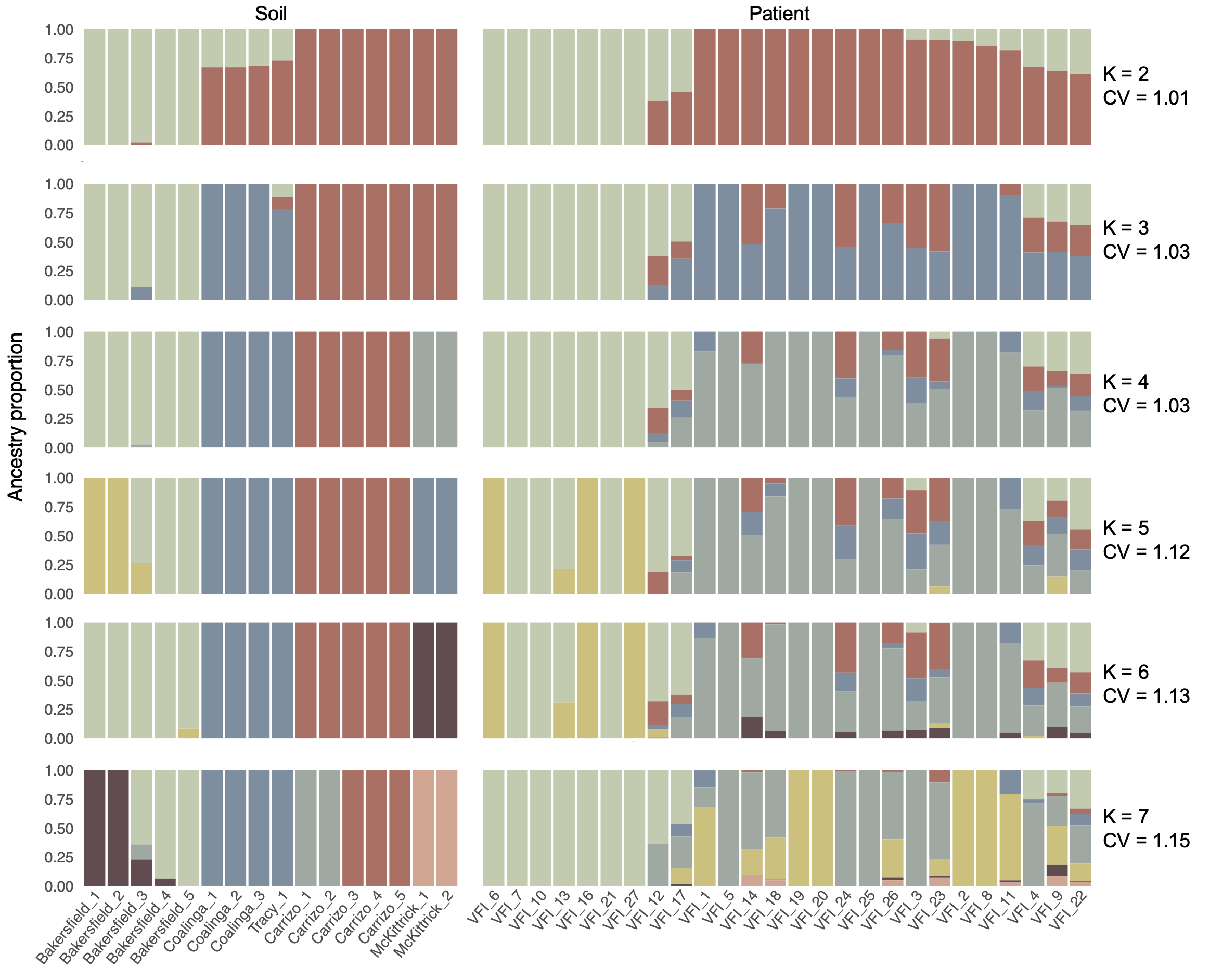

**Supplementary Figure S4.** Stacked bar plots showing ancestry proportions estimated by ADMIXTURE under *K* = 2–7 under the alternative specification. Herein, all isolates were included in the ADMIXTURE run, which included several clonal sets. Each bar represents an isolate, with soil-derived on the left and patient-derived on the right. The mean cross-validation error for a given *K* is listed on the right side of the plot.

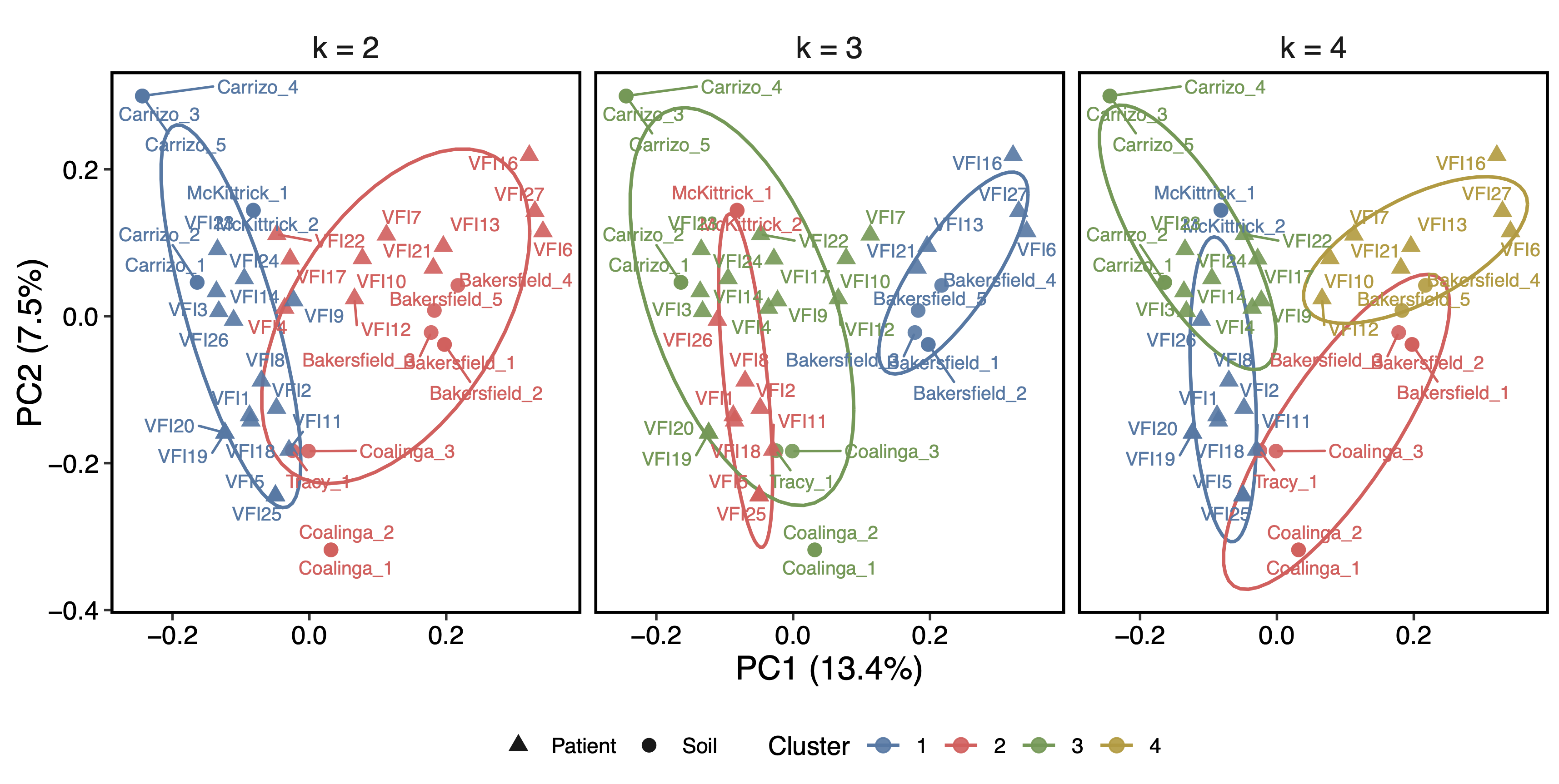

**Supplementary Figure S5.** PCA of the SNP matrix with K-means clustering. Soil-derived isolates (circles) and patient-derived isolates (triangles) are colored by their cluster assignment under K-means solutions of K = 2–4. Clustering was performed on coordinates in the first four principal components (32.6% of the total variance); only the first two PC axes are shown here. Ellipses are 68% confidence ellipses assuming bivariate normality within each cluster and are included as a visual aid. Note this method is distinct from ADMIXTURE clustering.

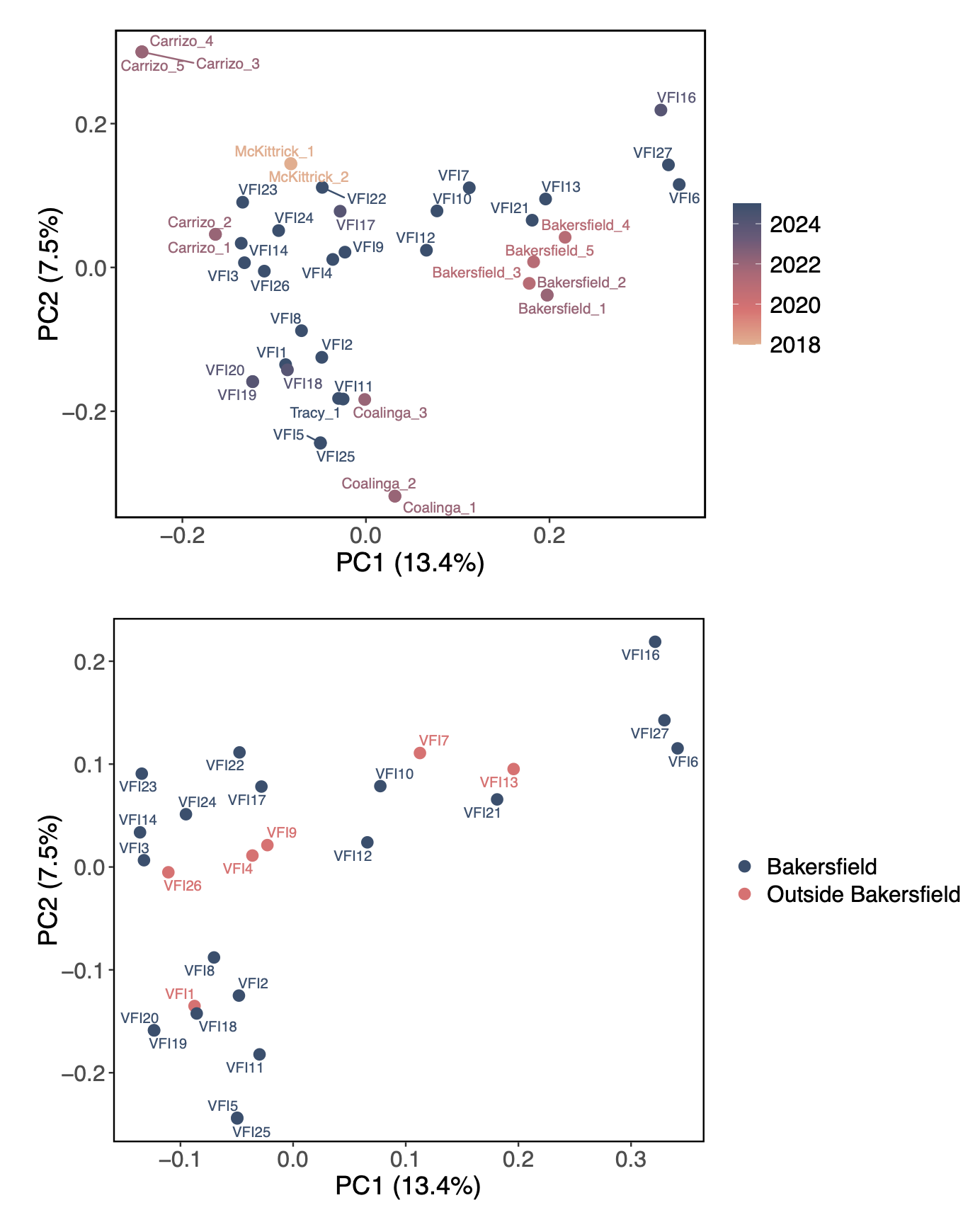

**Supplementary Figure S6.** PCA of the SNP matrix with isolates (points) colored by year of sample collection (top) or patient residence within or outside of Bakersfield zip codes (bottom). In the bottom plot, all isolates were included in the PCA, but only the patient isolates are plotted. The first two axes are shown with the percent variance explained in parentheses.

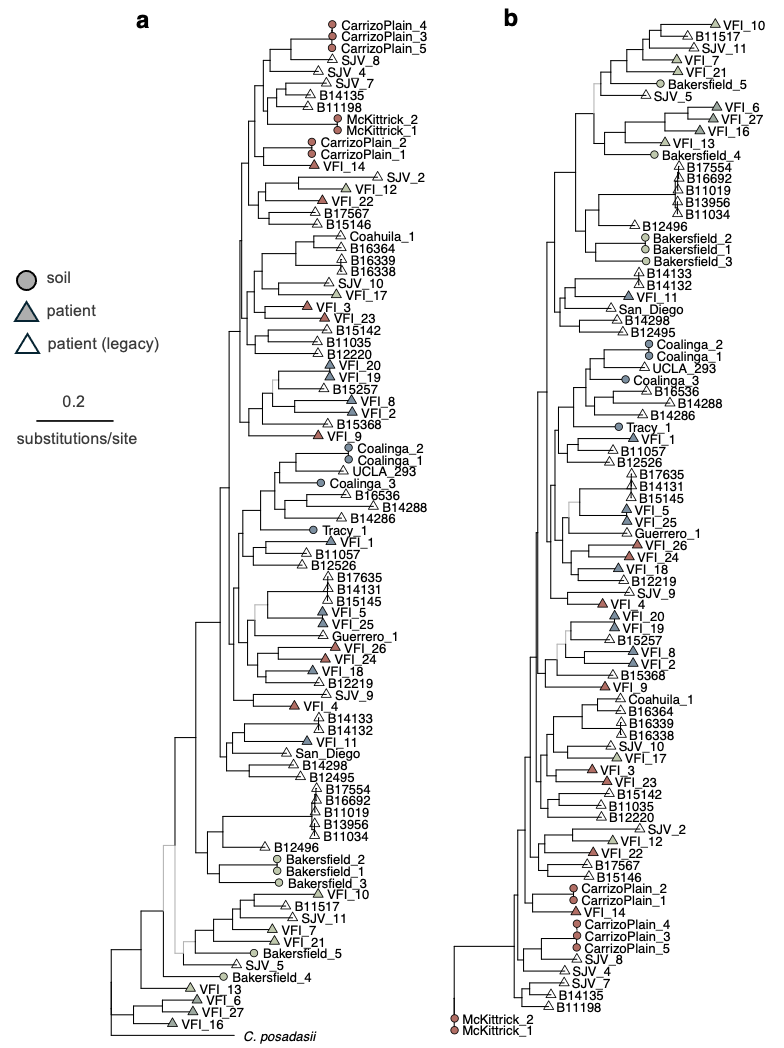

**Supplementary Figure S7.** Maximum likelihood phylogenies on the SNP matrix of all isolates including our soil (filled circles; n = 16) and patient (filled triangles; n = 26) samples and legacy patient genomes (open triangles; n = 44). Trees are either **(a)** rooted to *C. posadasii* (Silveira strain) or **(b)** unrooted. For both trees, branch lengths are proportional to the number of substitutions per site. Black branches are well supported, with SH-aLRT ≥ 80 and UFBoot ≥ 95 (i.e., the branch was produced in 95% of ultrafast bootstrap replicates); gray branches have low support. For both trees, tips are colored based on the dominant assignment from ADMIXTURE under *K* = 4. Legacy patient genomes were not included in the ADMXITURE run and are therefore unfilled.

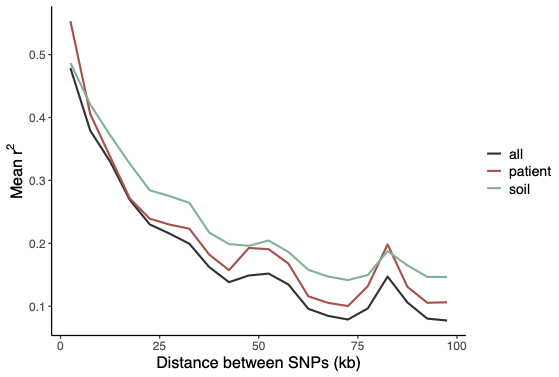

**Supplementary Figure S8.** Linkage disequilibrium (LD) decay curves for groups of our isolates (all, patient only, or soil only).

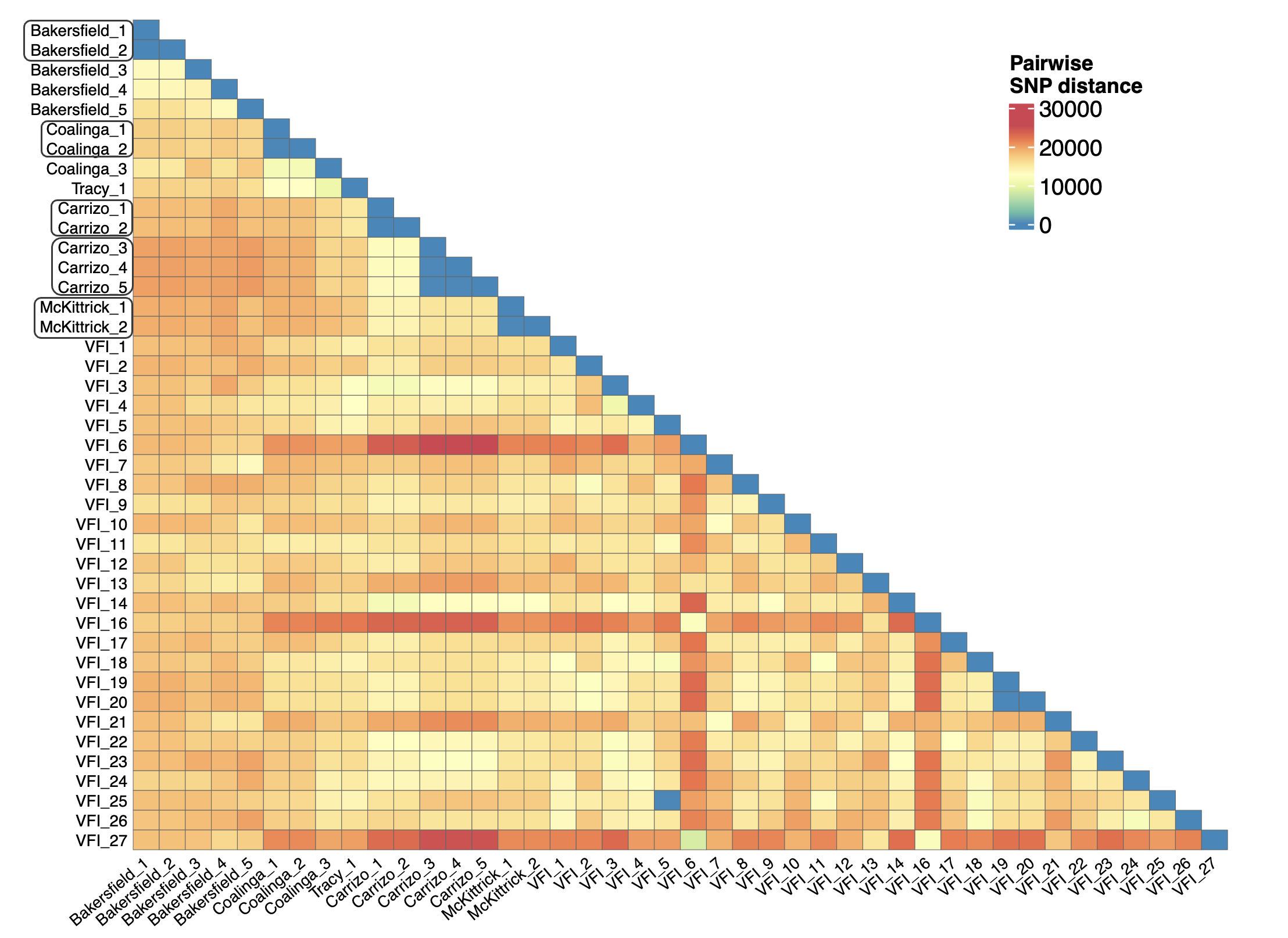

**Supplementary Figure S9.** Heatmap of pairwise SNP distances between all of our soil- (n = 16) and patient-derived (n = 26) genomes. Soil samples included in the same box on the left axis are those originating from the same rodent burrow. Note that VFI_5 and VFI_25 are *C. immitis* genomes originating from the same patient while all other patient-derived genomes are from distinct patients.

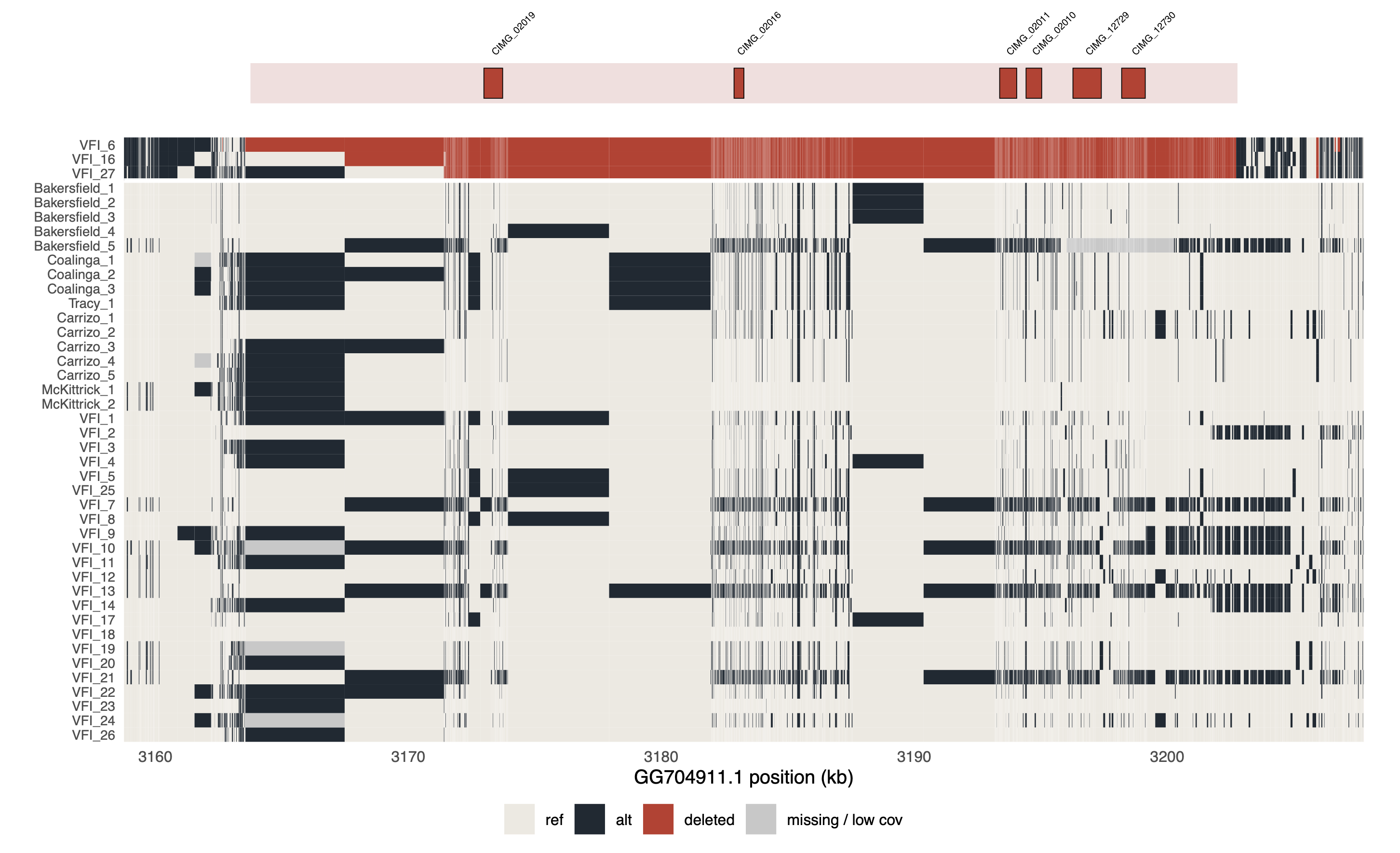

**Supplementary Figure S10.** Genotype matrix of the region surrounding the deletion shared by three patient-derived *C. immitis* isolates (VFI_6, VFI_16, and VFI_27). The top horizontal bar indicates the genomic positions and names of the six genes contained in the deleted region. Red in the genotype matrix heatmap denotes that no genotype was called at that position. We used genome coverage to determine that this region was deleted for the three patient isolates rather than simply not called.

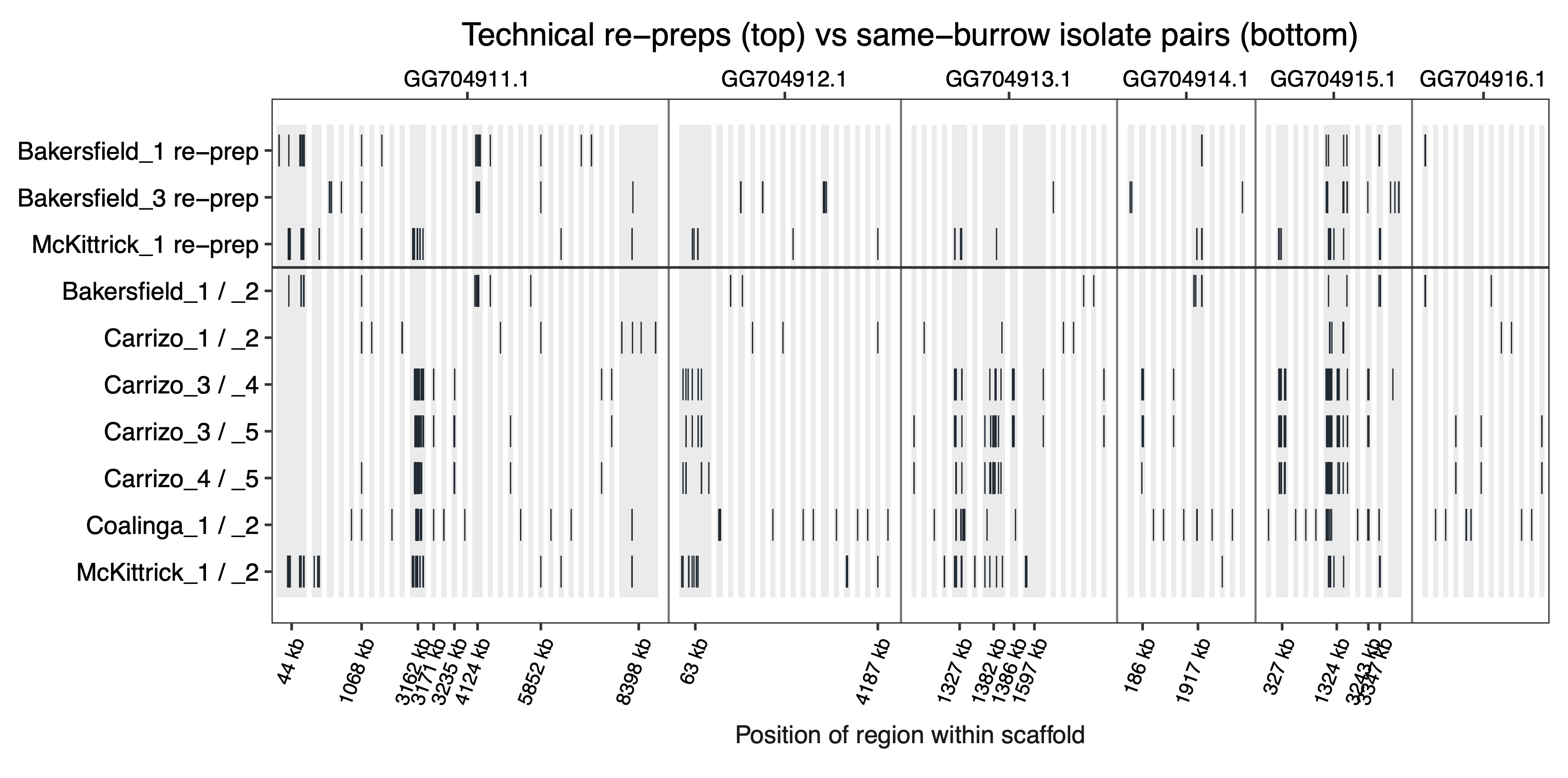

**Supplementary Figure S11.** Location of SNPs in technical replicates (top) and same-burrow pairs (bottom). SNPs are denoted with a vertical black line and shown based on their position within each scaffold. Intermediary sequences with no SNPs are removed.

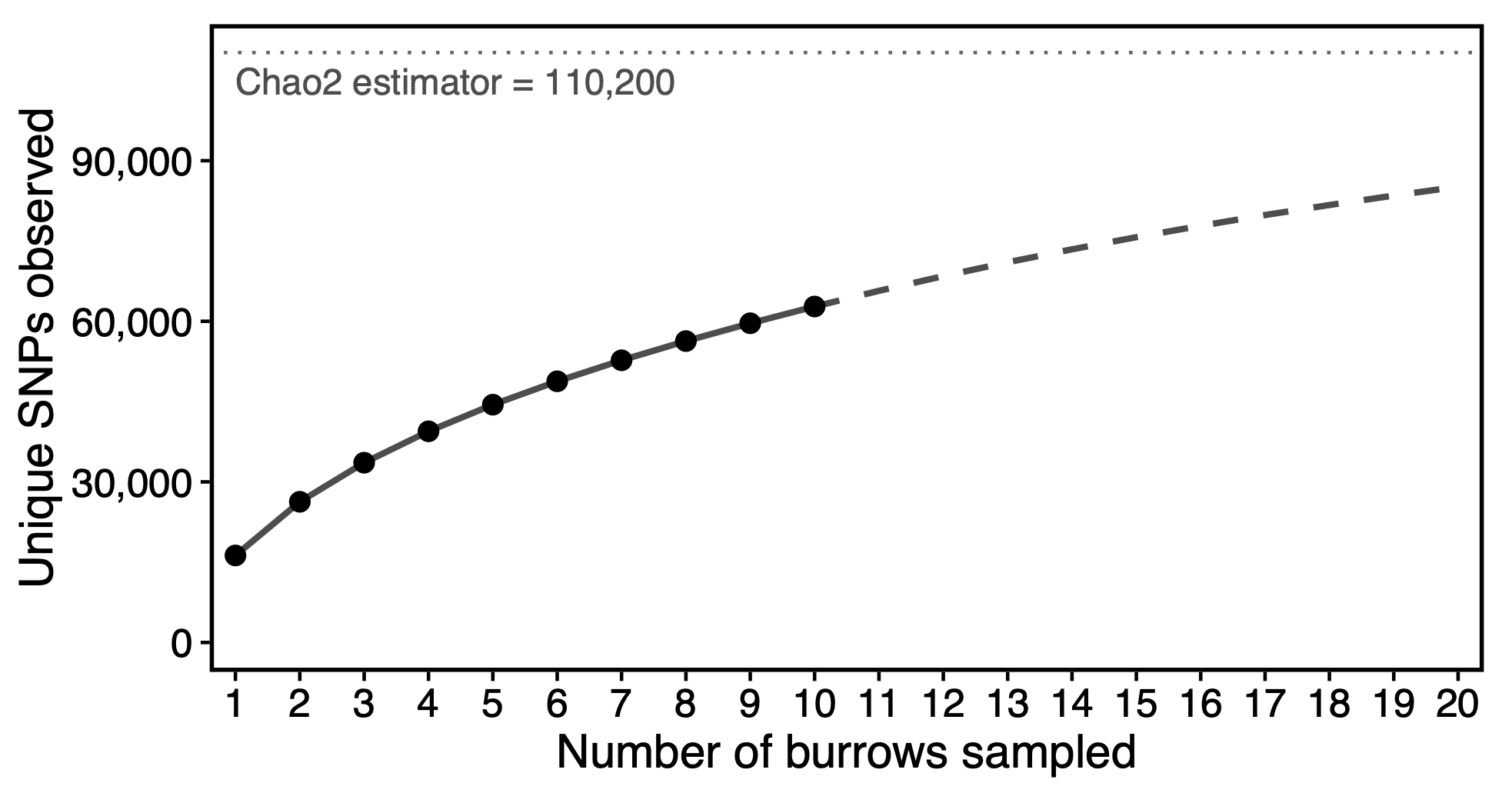

**Supplementary Figure S12.** SNP accumulation curve across burrows. The points and solid line denote the average number of unique SNPs observed for a given number of burrows. The dashed line extrapolates the SNP accumulation relationship from ten to twenty burrows following Colwell et al. 2012^23^. The asymptote of the curve (the Chao2 estimator) is denoted by the dotted horizontal line and was estimated at 110,200 SNPs.

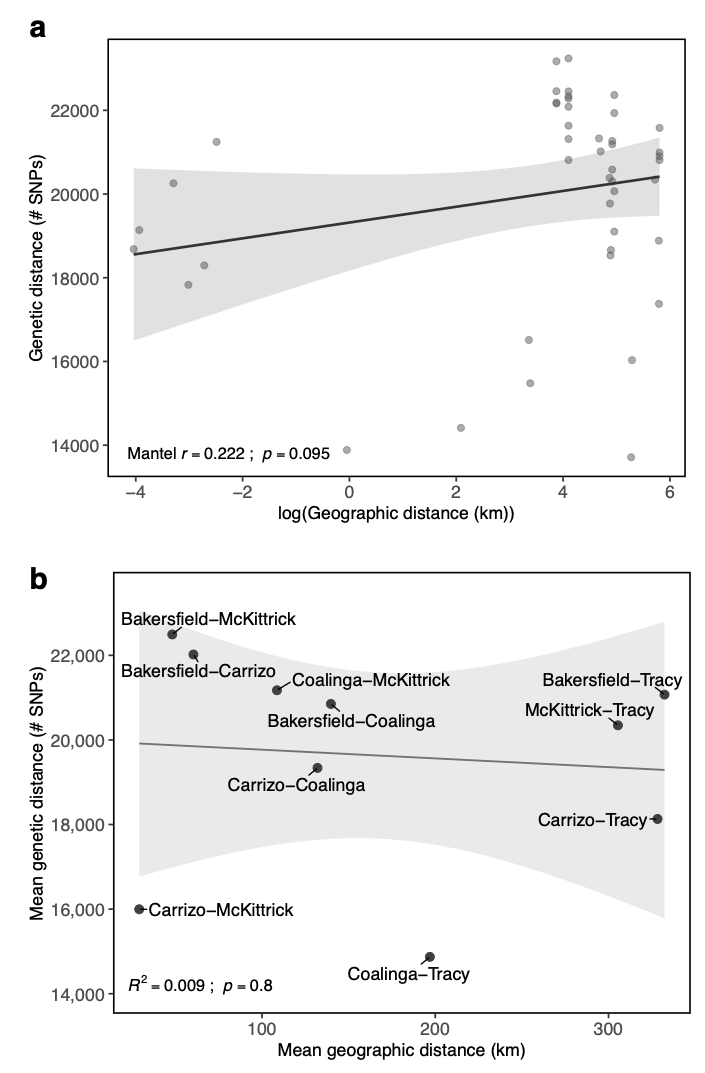

**Supplementary Figure S13.** (**a)** Scatterplot of genetic distance (here, the number of SNP sites that differ) against the log of geographic distance for pairs of soil isolates, including one representative isolate per burrow (n = 10 isolates, or 45 pairs). The Mantel test result is shown in the bottom left. Technical replicates were not included here to avoid violating assumptions of independence. **(b)** Scatterplot of genetic distance against geographic distance. Values for both distance measures are the mean between pairs of isolates from a given pair of sampling sites. The R^2^ and p-value from an ordinary least squares regression are shown in the bottom left.

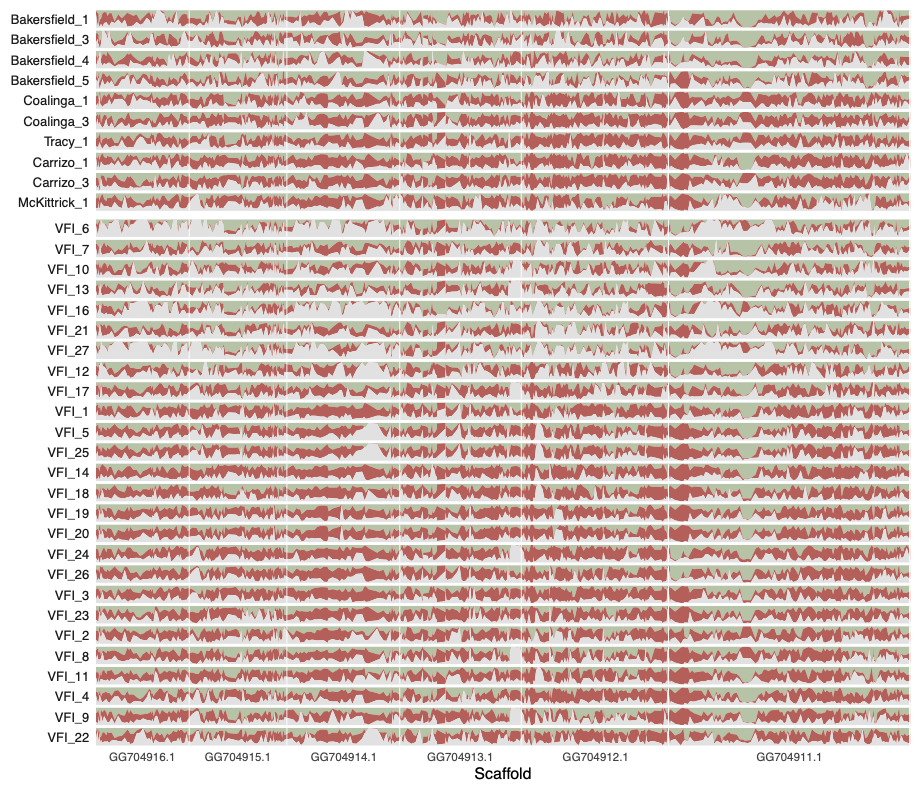

**Supplementary Figure S14.** Weightings for each of the three possible topologies in windows across the genome. Each horizontal bar corresponds to an isolate (soils on top; patients on bottom). The bar colors correspond to the topology branch colors with green, red, and white denoting placement in a clade with Bakersfield, non-Bakersfield, or its own branch, respectively. See Figure 4 in the main text for topology schematics.

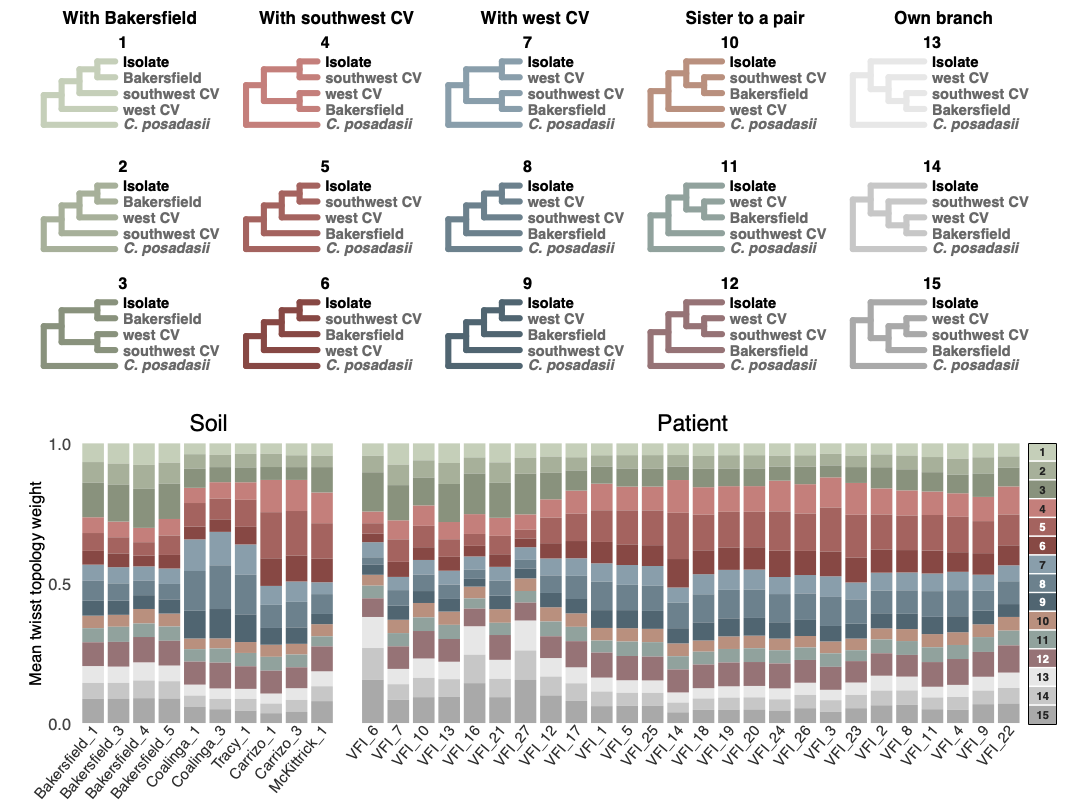

**Supplementary Figure S15.** (Top) All possible rooted topologies (n = 15) given three soil *C. immitis* reference populations, one focal isolate, and an outgroup (*C. posadasii*). (Bottom) Weightings for each topology, averaged across the genome, for each soil or patient isolate. The bar colors correspond to the topology branch colors in the top. Note that VFI_5 and VFI_25 are *C. immitis* isolates from the same patient while all other patient-derived isolates are from distinct patients.

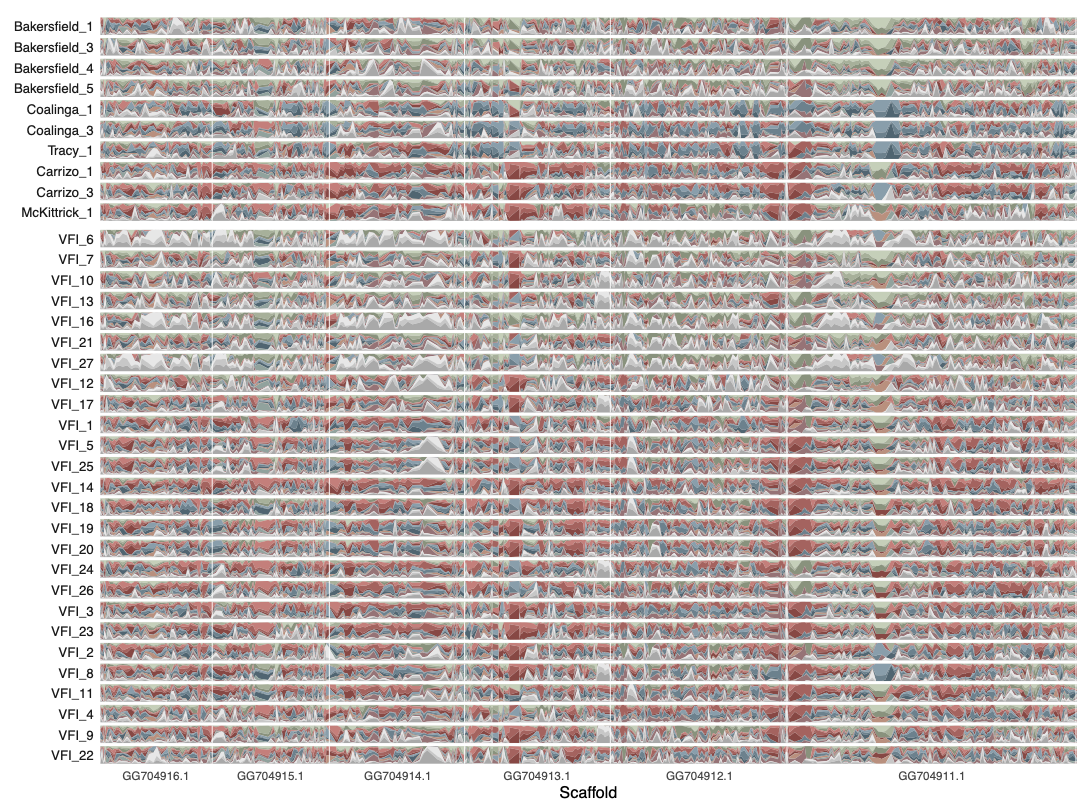

**Supplementary Figure S16.** Weightings for each of the 15 possible topologies in windows across the genome. Each horizontal bar corresponds to an isolate (soils on top; patients on bottom). The bar colors correspond to the topology branch colors denoted in Supplementary Figure S15. Note that VFI_5 and VFI_25 are *C. immitis* isolates from the same patient while all other patient-derived isolates are from distinct patients.

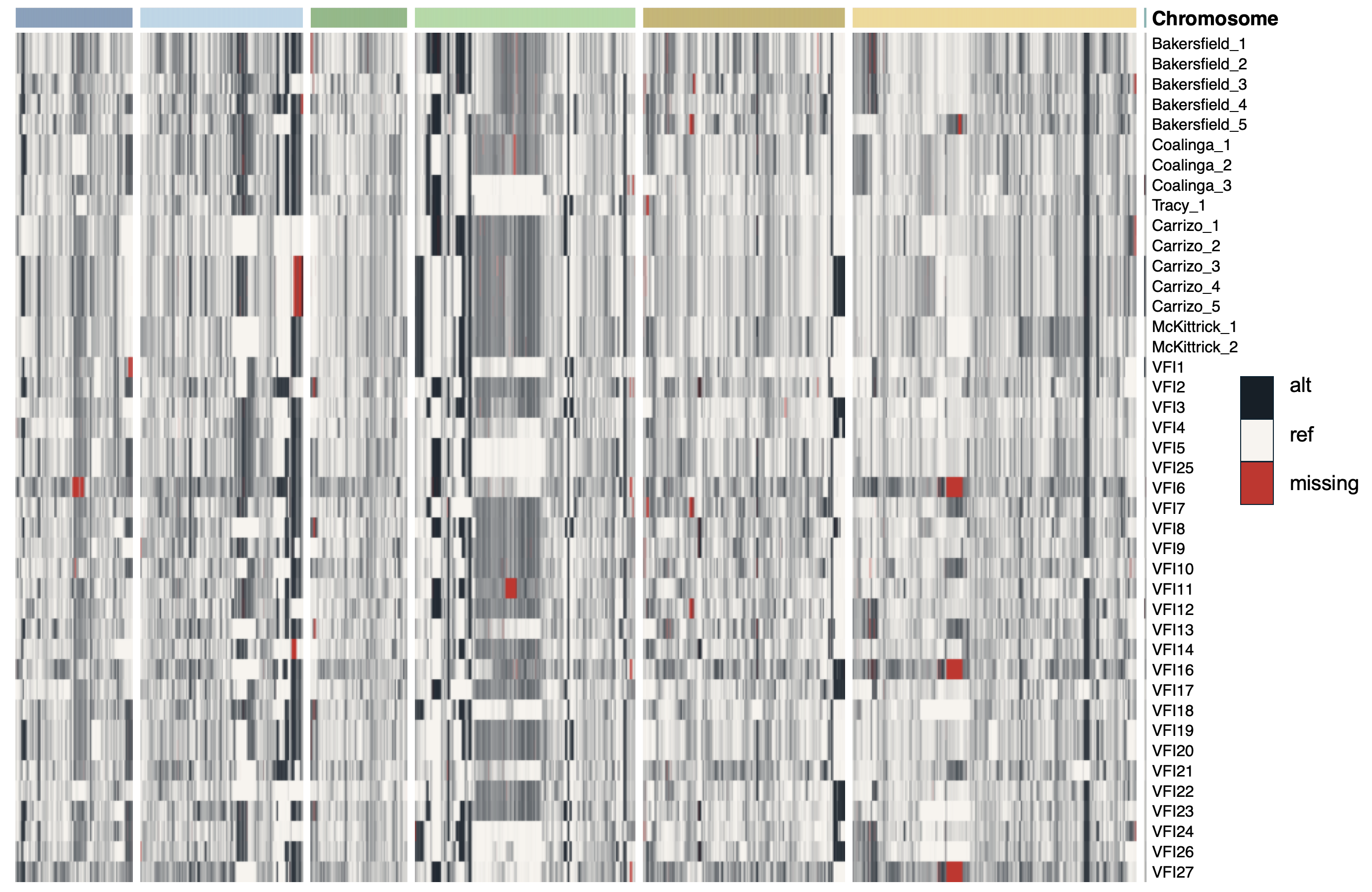

**Supplementary Figure S17.** Scaffold-level heatmap on the SNP matrix indicating alignment of soil- and patient-derived isolates (top 16 and bottom 26 samples, respectively) to the *C. immitis* reference genome (*C. immitis* RS GCF_000149335.2). Note that VFI_5 and VFI_25 are *C. immitis* isolates from the same patient while all other patient-derived isolates are from distinct patients. Off-white indicates sites with the same base call as the reference, black indicates an alternative call, and red indicates a missing call (~0.5% missingness across all samples). Scaffolds are plotted from left to right as: GG704916.1, GG704915.1, GG704914.1, GG704913.1, GG704912.1, GG704911.1, GG704917.1.

**Supplementary Tables**

**Supplementary Table S1.** Source metadata for *C. immitis* isolates generated in this study. Soil-derived isolates listed in a shared row originated from the same rodent burrow (but represent distinct colonies). Where reported, variation in precision of the latitude and longitude is due to differences in the GPS device used during field sampling. For patient-derived isolates the location denotes the clinic where patients were diagnosed, as their location of exposure is unknown.

|  | **Collection** | | | | |
| --- | --- | --- | --- | --- | --- |
| **Isolate** | **Year** | **Month** | **Location** | **Lat** | **Lon** |
| Bakersfield_1, Bakersfield_2 | 2022 | Jan | Kern River, Bakersfield, CA | 35.35577 | -119.10067 |
| Bakersfield_3 | 2021 | Jul | Kern River, Bakersfield, CA | 35.355441 | -119.10031 |
| Bakersfield_4 | 2021 | Oct | Kern River, Bakersfield, CA | 35.355401 | -119.10010 |
| Bakersfield_5 | 2021 | Apr | Kern River, Bakersfield, CA | 35.355342 | -119.09992 |
| Coalinga_1, Coalinga_2 | 2022 | Jan | Coalinga, CA | 36.158285 | -120.25092 |
| Coalinga_3 | 2022 | Jan | Coalinga, CA | 36.146602 | -120.33948 |
| Tracy_1 | 2025 | Jul | Tracy, CA | 37.640036 | -121.49113 |
| Carrizo_1, Carrizo_2 | 2022 | Apr | Carrizo Plain National Monument, Santa Margarita, CA | 35.0764545 | -119.6704473 |
| Carrizo_3, Carrizo_4, Carrizo_5 | 2022 | Apr | Carrizo Plain National Monument, Santa Margarita, CA | 35.0839818 | -119.6754326 |
| McKittrick_1, McKittrick_2 | 2018 | Mar | Highway 33 near McKittrick, CA | 35.339717 | -119.63083 |
| VFI_1 | 2025 | Jun | Valley Fever Institute at Kern Medical | NA | NA |
| VFI_2 | 2025 | Apr | Valley Fever Institute at Kern Medical | NA | NA |
| VFI_3 | 2025 | May | Valley Fever Institute at Kern Medical | NA | NA |
| VFI_4 | 2025 | Jul | Valley Fever Institute at Kern Medical | NA | NA |
| VFI_5 | 2025 | Jun | Valley Fever Institute at Kern Medical | NA | NA |
| VFI_6 | 2025 | Apr | Valley Fever Institute at Kern Medical | NA | NA |
| VFI_7 | 2025 | Jun | Valley Fever Institute at Kern Medical | NA | NA |
| VFI_8 | 2025 | Apr | Valley Fever Institute at Kern Medical | NA | NA |
| VFI_9 | 2025 | Feb | Valley Fever Institute at Kern Medical | NA | NA |
| VFI_10 | 2025 | Mar | Valley Fever Institute at Kern Medical | NA | NA |
| VFI_11 | 2025 | Mar | Valley Fever Institute at Kern Medical | NA | NA |
| VFI_12 | 2025 | Mar | Valley Fever Institute at Kern Medical | NA | NA |
| VFI_13 | 2025 | Mar | Valley Fever Institute at Kern Medical | NA | NA |
| VFI_14 | 2025 | Jan | Valley Fever Institute at Kern Medical | NA | NA |
| VFI_16 | 2024 | Nov | Valley Fever Institute at Kern Medical | NA | NA |
| VFI_17 | 2024 | Dec | Valley Fever Institute at Kern Medical | NA | NA |
| VFI_18 | 2024 | Oct | Valley Fever Institute at Kern Medical | NA | NA |
| VFI_19 | 2024 | Nov | Valley Fever Institute at Kern Medical | NA | NA |
| VFI_20 | 2024 | Nov | Valley Fever Institute at Kern Medical | NA | NA |
| VFI_21 | 2025 | Jul | Valley Fever Institute at Kern Medical | NA | NA |
| VFI_22 | 2025 | Oct | Valley Fever Institute at Kern Medical | NA | NA |
| VFI_23 | 2025 | Feb | Valley Fever Institute at Kern Medical | NA | NA |
| VFI_24 | 2025 | Feb | Valley Fever Institute at Kern Medical | NA | NA |
| VFI_25 | 2025 | Jun | Valley Fever Institute at Kern Medical | NA | NA |
| VFI_26 | 2025 | Sep | Valley Fever Institute at Kern Medical | NA | NA |
| VFI_27 | 2025 | Oct | Valley Fever Institute at Kern Medical | NA | NA |

**Supplementary Table S2.** Sequence metadata for all samples included in our analysis. ‘Year Sequenced’ refers to when sequences were generated (for the isolates generated in this study) or when sequences were published to the Sequence Read Archive (for previously generated isolates). All samples were sequenced on Illumina machines with paired-end reads.

| **Isolate** | **Biosample or SRA Accession** | **Year Sequenced** | **Sequencing Platform** | **Read Length (bp)** | **% mapped** | **% >10x coverage** |
| --- | --- | --- | --- | --- | --- | --- |
| Bakersfield_1 | SAMN62893303 | 2025 | NovaSeq 6000 | 150 | 88.09 | 97.60 |
| Bakersfield_2 | SAMN62893304 | 2025 | NovaSeq 6000 | 150 | 88.57 | 97.62 |
| Bakersfield_3 | SAMN62893305 | 2025 | NovaSeq 6000 | 150 | 86.91 | 97.59 |
| Bakersfield_4 | SAMN62893306 | 2025 | NovaSeq 6000 | 150 | 87.95 | 97.67 |
| Bakersfield_5 | SAMN62893307 | 2025 | NovaSeq 6000 | 150 | 81.59 | 98.56 |
| Coalinga_1 | SAMN62893308 | 2025 | NovaSeq 6000 | 150 | 88.13 | 98.23 |
| Coalinga_2 | SAMN62893309 | 2025 | NovaSeq 6000 | 150 | 87.54 | 98.23 |
| Coalinga_3 | SAMN62893310 | 2025 | NovaSeq 6000 | 150 | 87.47 | 98.65 |
| Tracy_1 | SAMN62893311 | 2025 | NovaSeq 6000 | 150 | 87.50 | 98.62 |
| Carrizo_1 | SAMN62893312 | 2025 | NovaSeq 6000 | 150 | 85.66 | 98.12 |
| Carrizo_2 | SAMN62893313 | 2025 | NovaSeq 6000 | 150 | 83.97 | 98.13 |
| Carrizo_3 | SAMN62893314 | 2024 | NovaSeq 6000 | 150 | 88.98 | 97.81 |
| Carrizo_4 | SAMN62893315 | 2024 | NovaSeq 6000 | 150 | 86.60 | 97.93 |
| Carrizo_5 | SAMN62893316 | 2024 | NovaSeq 6000 | 150 | 89.04 | 97.81 |
| McKittrick_1 | SAMN62893317 | 2025 | NovaSeq 6000 | 150 | 85.01 | 97.97 |
| McKittrick_2 | SAMN62893318 | 2025 | NovaSeq 6000 | 150 | 86.62 | 98.18 |
| VFI_1 | SAMN62893319 | 2026 | NovaSeq 6000 | 150 | 90.37 | 98.92 |
| VFI_2 | SAMN62893320 | 2026 | NovaSeq 6000 | 150 | 91.00 | 97.88 |
| VFI_3 | SAMN62893321 | 2026 | NovaSeq 6000 | 150 | 90.21 | 97.89 |
| VFI_4 | SAMN62893322 | 2026 | NovaSeq 6000 | 150 | 87.29 | 98.66 |
| VFI_5 | SAMN62893323 | 2026 | NovaSeq 6000 | 150 | 88.13 | 98.62 |
| VFI_6 | SAMN62893324 | 2026 | NovaSeq 6000 | 150 | 88.70 | 98.16 |
| VFI_7 | SAMN62893325 | 2026 | NovaSeq 6000 | 150 | 86.38 | 98.07 |
| VFI_8 | SAMN62893326 | 2026 | NovaSeq 6000 | 150 | 90.63 | 97.83 |
| VFI_9 | SAMN62893327 | 2026 | NovaSeq 6000 | 150 | 89.98 | 97.94 |
| VFI_10 | SAMN62893328 | 2026 | NovaSeq 6000 | 150 | 85.85 | 97.91 |
| VFI_11 | SAMN62893329 | 2026 | NovaSeq 6000 | 150 | 89.72 | 97.92 |
| VFI_12 | SAMN62893330 | 2026 | NovaSeq 6000 | 150 | 90.82 | 97.87 |
| VFI_13 | SAMN62893331 | 2026 | NovaSeq 6000 | 150 | 89.58 | 97.81 |
| VFI_14 | SAMN62893332 | 2026 | NovaSeq 6000 | 150 | 89.27 | 97.93 |
| VFI_16 | SAMN62893333 | 2026 | NovaSeq 6000 | 150 | 91.27 | 97.73 |
| VFI_17 | SAMN62893334 | 2026 | NovaSeq 6000 | 150 | 86.97 | 98.21 |
| VFI_18 | SAMN62893335 | 2026 | NovaSeq 6000 | 150 | 89.24 | 98.62 |
| VFI_19 | SAMN62893336 | 2026 | NovaSeq 6000 | 150 | 91.17 | 97.99 |
| VFI_20 | SAMN62893337 | 2026 | NovaSeq 6000 | 150 | 89.58 | 97.99 |
| VFI_21 | SAMN62893338 | 2026 | NovaSeq 6000 | 150 | 89.37 | 98.52 |
| VFI_22 | SAMN62893339 | 2026 | NovaSeq 6000 | 150 | 90.08 | 98.02 |
| VFI_23 | SAMN62893340 | 2026 | NovaSeq 6000 | 150 | 90.75 | 97.86 |
| VFI_24 | SAMN62893341 | 2026 | NovaSeq 6000 | 150 | 89.92 | 98.29 |
| VFI_25 | SAMN62893342 | 2026 | NovaSeq 6000 | 150 | 89.61 | 98.60 |
| VFI_26 | SAMN62893343 | 2026 | NovaSeq 6000 | 150 | 88.69 | 98.51 |
| VFI_27 | SAMN62893344 | 2026 | NovaSeq 6000 | 150 | 88.78 | 98.29 |
| B11019 | SRR8530936 | 2019 | HiSeq 2500 | 250 | 82.26 | 98.24 |
| B11034 | SRR8530937 | 2019 | HiSeq 2500 | 250 | 82.66 | 97.71 |
| B11035 | SRR8530933 | 2019 | HiSeq 2500 | 250 | 83.00 | 97.63 |
| B11057 | SRR21204687 | 2022 | HiSeq 2500 | 250 | 85.39 | 98.08 |
| B11198 | SRR8530934 | 2019 | HiSeq 2500 | 250 | 83.97 | 97.74 |
| B11517 | SRR8530935 | 2019 | HiSeq 2500 | 250 | 82.91 | 97.93 |
| B12219 | SRR21204724 | 2022 | HiSeq 2500 | 250 | 81.76 | 98.26 |
| B12220 | SRR21204723 | 2022 | HiSeq 2500 | 250 | 81.17 | 97.87 |
| B12495 | SRR7206601 | 2019 | HiSeq 2500 | 250 | 82.06 | 96.32 |
| B12496 | SRR7206598 | 2019 | HiSeq 2500 | 250 | 83.18 | 97.66 |
| B12526 | SRR21204713 | 2022 | HiSeq 2500 | 250 | 80.95 | 98.50 |
| B13956 | SRR7206599 | 2019 | HiSeq 2500 | 250 | 82.73 | 97.56 |
| B14131 | SRR21204705 | 2022 | HiSeq 2500 | 250 | 81.50 | 98.35 |
| B14132 | SRR21204704 | 2022 | HiSeq 2500 | 250 | 85.86 | 97.63 |
| B14133 | SRR21204703 | 2022 | HiSeq 2500 | 250 | 86.89 | 97.65 |
| B14135 | SRR21204701 | 2022 | HiSeq 2500 | 250 | 83.36 | 97.88 |
| B14286 | SRR21204700 | 2022 | HiSeq 2500 | 250 | 83.67 | 97.54 |
| B14288 | SRR21204699 | 2022 | HiSeq 2500 | 250 | 84.41 | 96.69 |
| B14298 | SRR21204695 | 2022 | HiSeq 2500 | 250 | 85.11 | 96.95 |
| B15142 | SRR21204694 | 2022 | HiSeq 2500 | 250 | 84.55 | 96.57 |
| B15145 | SRR21204693 | 2022 | HiSeq 2500 | 250 | 83.12 | 98.01 |
| B15146 | SRR21204692 | 2022 | HiSeq 2500 | 250 | 84.26 | 97.48 |
| B15257 | SRR21204691 | 2022 | HiSeq 2500 | 250 | 84.51 | 94.92 |
| B15368 | SRR21204689 | 2022 | HiSeq 2500 | 250 | 84.95 | 97.62 |
| B16338 | SRR21204684 | 2022 | HiSeq 2500 | 250 | 82.12 | 94.37 |
| B16339 | SRR21204683 | 2022 | HiSeq 2500 | 250 | 82.15 | 97.89 |
| B16364 | SRR21204682 | 2022 | HiSeq 2500 | 250 | 80.80 | 96.59 |
| B16536 | SRR21204680 | 2022 | HiSeq 2500 | 250 | 92.48 | 94.99 |
| B16692 | SRR21204679 | 2022 | HiSeq 2500 | 250 | 85.03 | 97.98 |
| B17554 | SRR21204677 | 2022 | HiSeq 2500 | 250 | 72.31 | 97.12 |
| B17567 | SRR21204675 | 2022 | HiSeq 2500 | 250 | 86.99 | 96.34 |
| B17635 | SRR21204674 | 2022 | HiSeq 2500 | 250 | 86.48 | 97.06 |
| Coahuila_1 | SRR3468021 | 2016 | HiSeq 2500 | 250 | 83.47 | 93.06 |
| San_Diego_1 | SRR3468019 | 2016 | HiSeq 2500 | 100 | 82.06 | 98.00 |
| SJV_10 | SRR3468081 | 2016 | HiSeq 2500 | 250 | 82.92 | 93.10 |
| SJV_11 | SRR3468017 | 2016 | HiSeq 2500 | 100 | 81.86 | 96.59 |
| SJV_2 | SRR3468016 | 2016 | HiSeq 2500 | 100 | 83.81 | 97.68 |
| SJV_4 | SRR3468038 | 2016 | HiSeq 2500 | 100 | 79.76 | 97.76 |
| SJV_5 | SRR3468049 | 2016 | HiSeq 2500 | 100 | 83.81 | 97.71 |
| SJV_7 | SRR3468071 | 2016 | HiSeq 2500 | 100 | 81.91 | 97.79 |
| SJV_8 | SRR3468079 | 2016 | HiSeq 2500 | 100 | 80.63 | 96.07 |
| SJV_9 | SRR3468080 | 2016 | HiSeq 2500 | 100 | 81.98 | 91.20 |
| UCLA_293 | SRR25626827 | 2023 | MiSeq | 250 | 87.43 | 90.92 |
| Guerrero_1 | SRR3468018 | 2016 | HiSeq 2500 | 100 | 82.55 | 96.28 |

**Supplementary Table S3.** Mean and weighted genome-wide F_ST_ (Weir and Cockerham) values between soil clusters inferred from ADMIXTURE with *K* = 2 or 3 (top) and *K* = 4 (bottom). The southwestern Central Valley cluster includes the Carrizo and McKittrick isolates; the western Central Valley cluster includes the Coalinga and Tracy isolates. We report sample sizes and F_ST_ estimates when including all isolates (left) and when excluding potential clones (i.e., those differing by <200 SNPs from another isolate).

|  |  | **All samples** | | | **Without near-clones** | | |
| --- | --- | --- | --- | --- | --- | --- | --- |
|  | **Soil cluster comparison** | **n** | **Mean F_ST_** | **Weighted F_ST_** | **n** | **Mean F_ST_** | **Weighted F_ST_** |
| *K* = 2 or 3 | Bakersfield vs non-Bakersfield | 5 v 11 | 0.162 | 0.241 | 4 v 6 | 0.073 | 0.146 |
| *K* = 4 | Bakersfield vs southwestern CV | 5 v 7 | 0.242 | 0.361 | 4 v 3 | 0.093 | 0.211 |
|  | Bakersfield vs western CV | 5 v 4 | 0.155 | 0.275 | 4 v 3 | 0.070 | 0.176 |
|  | Southwestern CV vs western CV | 7 v 4 | 0.282 | 0.394 | 3 v 3 | 0.112 | 0.234 |

**Supplementary Table S4.** Index of association ($\bar{r}_{d})$, initial r^2^ and half-decay distance (LD_50_) for our isolates (all, patient only, or soil only). The sample size here refers to the number of isolates included in the analysis when excluding potential clones.

|  | $\bar{r}_{d}$ | Initial r^2^ | LD_50_ |
| --- | --- | --- | --- |
| All (n = 34) | 0.011 | 0.479 | 22.5 kb |
| Patient only (n = 24) | 0.015 | 0.553 | 17.5 kb |
| Soil only (n = 10) | 0.013 | 0.487 | 37.5 kb |

**Supplementary Table S5.** Private alleles per burrow. We report the number and percent out of 62,759 total segregating SNPs among these isolates. We list the isolates associated with a given burrow (including only one of a given clonal pair).

| **Burrow** | **Isolates** | **# SNPs** | **% of total** |
| --- | --- | --- | --- |
| 1 | Bakersfield_1 | 4,051 | 6.46% |
| 2 | Bakersfield_3 | 4,048 | 6.45% |
| 3 | Bakersfield_4 | 4,009 | 6.39% |
| 4 | Bakersfield_5 | 4,551 | 7.25% |
| 5 | Coalinga_1 | 2,673 | 4.26% |
| 6 | Coalinga_3 | 2,214 | 3.53% |
| 7 | Tracy_1 | 2,543 | 4.05% |
| 8 | Carrizo_1 | 2,341 | 3.73% |
| 9 | Carrizo_3 | 3,104 | 4.95% |
| 10 | McKittrick_1 | 3,260 | 5.19% |

**Supplementary Table S6.** Geographic and genomic distances across scales. Scales include within-burrow, between-burrow at a given collection site, between collection sites. Further, we report pairwise distances between each of the five collection sites (Coalinga, Tracy, Carrizo, McKittrick, and Bakersfield). Variation among technical replicates, which are pairs known to be identical, is listed for context.

|  |  | **Geographic distance (km)** | | **Genomic distance (SNPs)** | |
| --- | --- | --- | --- | --- | --- |
|  | **n pairs** | **Mean** | **Range** | **Mean** | **Range (min, max)** |
| Technical replicates | 3 | NA | NA | 60 | 47–80 |
| Within-burrow | 7 | 0 | 0 | 105 | 29–169 |
| Between-burrow | 8 | 1.16 | 0.02–8.07 | 17,898 | 13,811–21,111 |
| Between-site (overall) | 37 | 142.78 | 28.76–332.35 | 20,300 | 13,627–23,126 |
| Coalinga-Tracy | 2 | 196.9 | 195.3–198.5 | 14,776 | 13,627–15,924 |
| Carrizo-McKittrick | 2 | 29.1 | 28.8–29.5 | 15,926 | 15,408–16,443 |
| Carrizo-Tracy | 2 | 328.3 | 327.8–328.7 | 18,038 | 17,259–18,816 |
| Carrizo-Coalinga | 4 | 132 | 130.4–133.6 | 19,199 | 18,419–20,247 |
| McKittrick-Tracy | 1 | 305.4 | 305.4 | 20,236 | 20,236 |
| Bakersfield-Coalinga | 8 | 139.8 | 137.0–142.5 | 20,743 | 18,963–22,226 |
| Bakersfield-Carrizo | 8 | 60.4 | 60.4–60.5 | 21,901 | 20,662–23,126 |
| Bakersfield-Tracy | 4 | 332.3 | 332.3 | 20,958 | 20,742–21,454 |
| Coalinga-McKittrick | 2 | 108.5 | 107.0–110.4 | 21,038 | 20,875–21,202 |
| Bakersfield-McKittrick | 4 | 48.2 | 48.2 | 22,383 | 22,069–23,048 |

**Supplementary Table S7.** Diversity metrics for sets of genomes. Technical replicates and near-clones (i.e., those differing by <200 SNPs from another isolate) were excluded from these calculations. Metrics were calculated using a shared VCF filtered to remove variants with a minor allele frequency < 0.05, thus causing downward bias in the reported diversity metrics. Metrics were normalized by the number of callable sites. S refers to the number of segregating sites; θ_w_ refers to Watterson’s estimator; π refers to nucleotide diversity.

|  | **Total # genomes** | **# genomes**  **(no near-clones)** | **S** | **θ_w_** | **π** |
| --- | --- | --- | --- | --- | --- |
| All soil | 16 | 10 | 43,129 | 6.46 x 10^-4^ | 6.34 x 10^-4^ |
| All patient | 26 | 24 | 50,901 | 5.77 x 10^-4^ | 6.58 x 10^-4^ |
| All soil and patient | 42 | 34 | 53,535 | 5.55 x 10^-4^ | 6.70 x 10^-4^ |
| All samples (with legacy) | 86 | 78 | 55,874 | 4.80 x 10^-4^ | 6.73 x 10^-4^ |

**Supplementary Table S8**. Mating type loci assignment for all isolates. Soil isolates originating from the same rodent burrow are noted with matching superscript letters.

| **Isolate** | **Source** | **Mating type assignment** |
| --- | --- | --- |
| Bakersfield_1^a^ | Soil | MAT1-2 |
| Bakersfield_2^a^ | Soil | MAT1-2 |
| Bakersfield_3 | Soil | MAT1-1 |
| Bakersfield_4 | Soil | MAT1-1 |
| Bakersfield_5 | Soil | MAT1-1 |
| Coalinga_1^e^ | Soil | MAT1-2 |
| Coalinga_2^e^ | Soil | MAT1-2 |
| Coalinga_3 | Soil | MAT1-2 |
| Tracy_1 | Soil | MAT1-1 |
| Carrizo_1^b^ | Soil | MAT1-1 |
| Carrizo_2^b^ | Soil | MAT1-1 |
| Carrizo_3^c^ | Soil | MAT1-1 |
| Carrizo_4^c^ | Soil | MAT1-1 |
| Carrizo_5^c^ | Soil | MAT1-1 |
| McKittrick_1^d^ | Soil | MAT1-1 |
| McKittrick_2^d^ | Soil | MAT1-1 |
| VFI_1 | Patient | MAT1-1 |
| VFI_2 | Patient | MAT1-1 |
| VFI_3 | Patient | MAT1-2 |
| VFI_4 | Patient | MAT1-2 |
| VFI_5 | Patient | MAT1-2 |
| VFI_6 | Patient | MAT1-2 |
| VFI_7 | Patient | MAT1-2 |
| VFI_8 | Patient | MAT1-2 |
| VFI_9 | Patient | MAT1-2 |
| VFI_10 | Patient | MAT1-1 |
| VFI_11 | Patient | MAT1-2 |
| VFI_12 | Patient | MAT1-1 |
| VFI_13 | Patient | MAT1-1 |
| VFI_14 | Patient | MAT1-1 |
| VFI_16 | Patient | MAT1-2 |
| VFI_17 | Patient | MAT1-1 |
| VFI_18 | Patient | MAT1-2 |
| VFI_19 | Patient | MAT1-1 |
| VFI_20 | Patient | MAT1-1 |
| VFI_21 | Patient | MAT1-1 |
| VFI_22 | Patient | MAT1-2 |
| VFI_23 | Patient | MAT1-2 |
| VFI_24 | Patient | MAT1-1 |
| VFI_25 | Patient | MAT1-2 |
| VFI_26 | Patient | MAT1-2 |
| VFI_27 | Patient | MAT1-1 |

**Supplementary Table S9.** Results from topology weighting by iterative sampling of subtrees (‘*Twisst*’) under two soil reference groups. For each isolate, we report the most common topology estimated by *Twisst* and its corresponding nearest grouping (‘Bak’, ‘non-Bak’ and ‘own’ refers to ‘Bakersfield’, ‘non-Bakersfield’, and ‘own branch’, respectively). We use one representative isolate per burrow, given the observed clonality. See Figure 4 for a schematic of the three possible topologies. We also report the proportion of windows where the modal topology was dominant, and the proportion of windows where the isolate was classified in its own branch. Values denote the mean and 95% confidence interval from the 100 bootstrap replicates.

|  |  |  | **Prop. modal topology** | | **Prop. own branch** | |
| --- | --- | --- | --- | --- | --- | --- |
| **Isolate** | **Source** | **Modal Topology** | **Mean** | **95% CI** | **Mean** | **95% CI** |
| Bakersfield_1  (clonal with Bakersfield_2) | Soil | Bak | 0.409 | [0.385, 0.440] | 0.299 | [0.276, 0.329] |
| Bakersfield_3 | Soil | Bak | 0.443 | [0.421, 0.467] | 0.303 | [0.280, 0.328] |
| Bakersfield_4 | Soil | Bak | 0.464 | [0.437, 0.491] | 0.303 | [0.273, 0.331] |
| Bakersfield_5 | Soil | Bak | 0.425 | [0.399, 0.461] | 0.309 | [0.281, 0.337] |
| Coalinga_1 | Soil | non-Bak | 0.507 | [0.481, 0.532] | 0.251 | [0.223, 0.280] |
| Coalinga_3 | Soil | non-Bak | 0.546 | [0.515, 0.575] | 0.229 | [0.204, 0.264] |
| Tracy_1 | Soil | non-Bak | 0.547 | [0.519, 0.578] | 0.211 | [0.184, 0.236] |
| Carrizo_1  (clonal with Carrizo_2) | Soil | non-Bak | 0.585 | [0.555, 0.613] | 0.217 | [0.190, 0.251] |
| Carrizo_3  (clonal with Carrizo_4 and 5) | Soil | non-Bak | 0.568 | [0.540, 0.592] | 0.227 | [0.205, 0.250] |
| McKittrick_1  (clonal with McKittrick_2) | Soil | non-Bak | 0.466 | [0.441, 0.495] | 0.308 | [0.284, 0.331] |
| VFI_1 | Patient | non-Bak | 0.541 | [0.505, 0.573] | 0.269 | [0.242, 0.296] |
| VFI_2 | Patient | non-Bak | 0.512 | [0.482, 0.535] | 0.278 | [0.254, 0.303] |
| VFI_3 | Patient | non-Bak | 0.611 | [0.588, 0.634] | 0.229 | [0.206, 0.255] |
| VFI_4 | Patient | non-Bak | 0.511 | [0.482, 0.543] | 0.236 | [0.210, 0.268] |
| VFI_5 | Patient | non-Bak | 0.515 | [0.489, 0.550] | 0.268 | [0.237, 0.293] |
| VFI_6 | Patient | own | 0.429 | [0.403, 0.454] | 0.512 | [0.482, 0.549] |
| VFI_7 | Patient | Bak | 0.446 | [0.415, 0.477] | 0.303 | [0.276, 0.331] |
| VFI_8 | Patient | non-Bak | 0.489 | [0.460, 0.513] | 0.278 | [0.255, 0.306] |
| VFI_9 | Patient | non-Bak | 0.464 | [0.436, 0.491] | 0.287 | [0.254, 0.313] |
| VFI_10 | Patient | non-Bak | 0.378 | [0.361, 0.403] | 0.351 | [0.324, 0.378] |
| VFI_11 | Patient | non-Bak | 0.522 | [0.493, 0.544] | 0.235 | [0.208, 0.266] |
| VFI_12 | Patient | non-Bak | 0.383 | [0.362, 0.409] | 0.338 | [0.310, 0.368] |
| VFI_13 | Patient | Bak | 0.454 | [0.423, 0.483] | 0.326 | [0.296, 0.352] |
| VFI_14 | Patient | non-Bak | 0.611 | [0.581, 0.640] | 0.211 | [0.184, 0.244] |
| VFI_16 | Patient | own | 0.401 | [0.384, 0.423] | 0.475 | [0.447, 0.505] |
| VFI_17 | Patient | non-Bak | 0.473 | [0.441, 0.502] | 0.307 | [0.279, 0.337] |
| VFI_18 | Patient | non-Bak | 0.553 | [0.517, 0.577] | 0.236 | [0.212, 0.259] |
| VFI_19 | Patient | non-Bak | 0.551 | [0.520, 0.580] | 0.247 | [0.220, 0.273] |
| VFI_20 | Patient | non-Bak | 0.550 | [0.519, 0.582] | 0.249 | [0.223, 0.274] |
| VFI_21 | Patient | Bak | 0.417 | [0.387, 0.447] | 0.338 | [0.307, 0.371] |
| VFI_22 | Patient | non-Bak | 0.513 | [0.485, 0.544] | 0.296 | [0.269, 0.318] |
| VFI_23 | Patient | non-Bak | 0.559 | [0.534, 0.585] | 0.262 | [0.235, 0.290] |
| VFI_24 | Patient | non-Bak | 0.587 | [0.562, 0.617] | 0.217 | [0.194, 0.243] |
| VFI_25 | Patient | non-Bak | 0.515 | [0.487, 0.548] | 0.267 | [0.236, 0.292] |
| VFI_26 | Patient | non-Bak | 0.545 | [0.516, 0.573] | 0.25 | [0.226, 0.279] |
| VFI_27 | Patient | own | 0.417 | [0.402, 0.438] | 0.493 | [0.467, 0.525] |

**Supplementary Table S10.** Results from topology weighting by iterative sampling of subtrees (‘*Twisst*’) under two soil reference groups (three possible topologies). We use one representative isolate per burrow, given the observed clonality. For each isolate, we report the rate at which the dominant topology switched between adjacent windows, and the average run of the dominant topology in units of number of windows and kb. Values denote the mean and 95% confidence interval from the 100 bootstrap replicates.

|  |  | **Switch rate** | | **Runs (in windows)** | | **Runs (in kb)** | |
| --- | --- | --- | --- | --- | --- | --- | --- |
| **Isolate** | **Source** | **Mean** | **95% CI** | **Mean** | **95% CI** | **Mean** | **95% CI** |
| Bakersfield_1  (clonal with Bakersfield_2) | Soil | 0.531 | [0.499, 0.565] | 1.86 | [1.75, 1.97] | 108.4 | [101.6, 115.3] |
| Bakersfield_3 | Soil | 0.526 | [0.489, 0.561] | 1.88 | [1.76, 2.01] | 109.2 | [102.1, 117.7] |
| Bakersfield_4 | Soil | 0.551 | [0.505, 0.590] | 1.80 | [1.68, 1.95] | 104.2 | [97.1, 112.7] |
| Bakersfield_5 | Soil | 0.563 | [0.523, 0.605] | 1.76 | [1.64, 1.89] | 101.3 | [94.6, 109.4] |
| Coalinga_1 | Soil | 0.535 | [0.499, 0.568] | 1.85 | [1.74, 1.98] | 106.4 | [100.0, 114.6] |
| Coalinga_3 | Soil | 0.521 | [0.482, 0.552] | 1.89 | [1.79, 2.04] | 109.1 | [102.6, 118.0] |
| Tracy_1 | Soil | 0.519 | [0.483, 0.557] | 1.90 | [1.77, 2.04] | 110.0 | [101.8, 117.8] |
| Carrizo_1  (clonal with Carrizo_2) | Soil | 0.481 | [0.448, 0.524] | 2.05 | [1.88, 2.19] | 118.3 | [108.1, 126.6] |
| Carrizo_3  (clonal with Carrizo_4 and 5) | Soil | 0.484 | [0.451, 0.527] | 2.04 | [1.87, 2.18] | 117.9 | [108.5, 126.0] |
| McKittrick_1  (clonal with McKittrick_2) | Soil | 0.505 | [0.472, 0.549] | 1.96 | [1.80, 2.08] | 113.1 | [103.6, 120.4] |
| VFI_1 | Patient | 0.502 | [0.465, 0.530] | 1.97 | [1.86, 2.11] | 113.4 | [107.1, 122.2] |
| VFI_2 | Patient | 0.518 | [0.488, 0.554] | 1.91 | [1.78, 2.02] | 110.4 | [102.6, 118.0] |
| VFI_3 | Patient | 0.466 | [0.429, 0.502] | 2.12 | [1.96, 2.28] | 122.5 | [113.6, 132.5] |
| VFI_4 | Patient | 0.501 | [0.459, 0.537] | 1.97 | [1.84, 2.14] | 113.6 | [104.9, 124.6] |
| VFI_5 | Patient | 0.530 | [0.498, 0.574] | 1.86 | [1.72, 1.98] | 107.8 | [99.8, 114.5] |
| VFI_6 | Patient | 0.485 | [0.451, 0.517] | 2.03 | [1.91, 2.18] | 117.4 | [110.1, 125.5] |
| VFI_7 | Patient | 0.535 | [0.499, 0.566] | 1.85 | [1.75, 1.97] | 107.3 | [100.4, 114.5] |
| VFI_8 | Patient | 0.517 | [0.478, 0.552] | 1.91 | [1.79, 2.06] | 110.3 | [103.1, 119.6] |
| VFI_9 | Patient | 0.533 | [0.497, 0.572] | 1.85 | [1.73, 1.98] | 107.6 | [99.9, 114.8] |
| VFI_10 | Patient | 0.555 | [0.518, 0.585] | 1.78 | [1.69, 1.91] | 102.4 | [96.8, 110.4] |
| VFI_11 | Patient | 0.509 | [0.471, 0.540] | 1.94 | [1.83, 2.09] | 112.3 | [105.2, 120.8] |
| VFI_12 | Patient | 0.538 | [0.507, 0.569] | 1.84 | [1.74, 1.94] | 105.8 | [100.1, 112.7] |
| VFI_13 | Patient | 0.561 | [0.532, 0.601] | 1.76 | [1.65, 1.85] | 102.0 | [94.8, 107.4] |
| VFI_14 | Patient | 0.473 | [0.439, 0.515] | 2.08 | [1.91, 2.24] | 120.7 | [111.0, 130.1] |
| VFI_16 | Patient | 0.455 | [0.421, 0.488] | 2.16 | [2.02, 2.33] | 125.7 | [117.3, 135.3] |
| VFI_17 | Patient | 0.534 | [0.502, 0.574] | 1.85 | [1.72, 1.96] | 106.9 | [99.8, 113.7] |
| VFI_18 | Patient | 0.518 | [0.479, 0.552] | 1.91 | [1.79, 2.06] | 110.6 | [103.9, 119.7] |
| VFI_19 | Patient | 0.528 | [0.485, 0.563] | 1.87 | [1.76, 2.03] | 108.4 | [101.1, 117.6] |
| VFI_20 | Patient | 0.527 | [0.483, 0.564] | 1.87 | [1.75, 2.04] | 108.5 | [100.5, 118.0] |
| VFI_21 | Patient | 0.552 | [0.513, 0.580] | 1.79 | [1.71, 1.92] | 103.1 | [97.7, 110.4] |
| VFI_22 | Patient | 0.535 | [0.494, 0.578] | 1.85 | [1.71, 2.00] | 107.0 | [98.9, 116.3] |
| VFI_23 | Patient | 0.495 | [0.466, 0.526] | 1.99 | [1.88, 2.11] | 115.4 | [108.2, 122.7] |
| VFI_24 | Patient | 0.466 | [0.430, 0.501] | 2.11 | [1.97, 2.28] | 121.9 | [112.6, 132.4] |
| VFI_25 | Patient | 0.530 | [0.497, 0.567] | 1.87 | [1.74, 1.98] | 107.9 | [100.5, 114.5] |
| VFI_26 | Patient | 0.502 | [0.467, 0.535] | 1.97 | [1.84, 2.11] | 113.4 | [105.7, 121.7] |
| VFI_27 | Patient | 0.457 | [0.425, 0.490] | 2.15 | [2.01, 2.31] | 124.6 | [116.0, 133.2] |

**Supplementary Table S11.** Results from topology weighting by iterative sampling of subtrees (‘*Twisst*’) under three soil reference groups. For each isolate, we report the most common topology estimated by *Twisst* and its corresponding nearest grouping (‘Bak’, ‘wCV’ , ‘swCV’ and ‘own’ refers to ‘Bakersfield’, ‘western Central Valley’, ‘southwestern Central Valley’, and ‘own branch’, respectively). The western Central Valley group includes the isolates from Coalinga and Tracy. We use one representative isolate per burrow, given the observed clonality. See Supplementary Figure S15 for a schematic of the 15 possible topologies and their corresponding numbers. We also report the proportion of windows where the modal topology was dominant, and the proportion of windows where the isolate was classified in its own branch.

|  |  |  | **Prop. modal topology** | | **Prop. own branch** | |
| --- | --- | --- | --- | --- | --- | --- |
| **Isolate** | **Source** | **Modal Topology** | **Mean** | **95% CI** | **Mean** | **95% CI** |
| Bakersfield_1  (clonal with Bakersfield_2) | Soil | Bak | 0.138 | [0.112, 0.158] | 0.216 | [0.187, 0.243] |
| Bakersfield_3 | Soil | Bak | 0.159 | [0.140, 0.180] | 0.198 | [0.179, 0.222] |
| Bakersfield_4 | Soil | Bak | 0.162 | [0.134, 0.187] | 0.229 | [0.206, 0.258] |
| Bakersfield_5 | Soil | Bak | 0.150 | [0.130, 0.175] | 0.225 | [0.194, 0.253] |
| Coalinga_1 | Soil | wCV | 0.190 | [0.167, 0.214] | 0.145 | [0.123, 0.167] |
| Coalinga_3 | Soil | wCV | 0.210 | [0.189, 0.245] | 0.132 | [0.107, 0.155] |
| Tracy_1 | Soil | wCV | 0.181 | [0.161, 0.206] | 0.128 | [0.106, 0.152] |
| Carrizo_1  (clonal with Carrizo_2) | Soil | swCV | 0.217 | [0.193, 0.240] | 0.113 | [0.092, 0.138] |
| Carrizo_3  (clonal with Carrizo_4 and 5) | Soil | swCV | 0.205 | [0.180, 0.232] | 0.138 | [0.116, 0.155] |
| McKittrick_1  (clonal with McKittrick_2) | Soil | swCV | 0.153 | [0.137, 0.171] | 0.207 | [0.184, 0.230] |
| VFI_1 | Patient | swCV | 0.139 | [0.120, 0.160] | 0.174 | [0.153, 0.199] |
| VFI_2 | Patient | swCV | 0.159 | [0.134, 0.183] | 0.182 | [0.162, 0.204] |
| VFI_3 | Patient | swCV | 0.199 | [0.176, 0.226] | 0.124 | [0.108, 0.145] |
| VFI_4 | Patient | swCV | 0.158 | [0.132, 0.177] | 0.146 | [0.120, 0.167] |
| VFI_5 | Patient | swCV | 0.147 | [0.122, 0.170] | 0.172 | [0.154, 0.194] |
| VFI_6 | Patient | Bak | 0.173 | [0.155, 0.197] | 0.437 | [0.410, 0.467] |
| VFI_7 | Patient | Bak | 0.149 | [0.125, 0.174] | 0.223 | [0.199, 0.245] |
| VFI_8 | Patient | swCV | 0.144 | [0.124, 0.166] | 0.184 | [0.166, 0.204] |
| VFI_9 | Patient | swCV | 0.132 | [0.111, 0.148] | 0.190 | [0.167, 0.214] |
| VFI_10 | Patient | Bak | 0.121 | [0.101, 0.141] | 0.256 | [0.230, 0.281] |
| VFI_11 | Patient | swCV | 0.157 | [0.132, 0.182] | 0.142 | [0.122, 0.163] |
| VFI_12 | Patient | swCV | 0.118 | [0.098, 0.135] | 0.253 | [0.224, 0.281] |
| VFI_13 | Patient | Bak | 0.167 | [0.148, 0.188] | 0.246 | [0.222, 0.267] |
| VFI_14 | Patient | swCV | 0.206 | [0.184, 0.231] | 0.110 | [0.091, 0.129] |
| VFI_16 | Patient | Bak | 0.179 | [0.157, 0.199] | 0.399 | [0.376, 0.427] |
| VFI_17 | Patient | swCV | 0.115 | [0.099, 0.137] | 0.231 | [0.205, 0.252] |
| VFI_18 | Patient | swCV | 0.154 | [0.132, 0.179] | 0.121 | [0.101, 0.144] |
| VFI_19 | Patient | swCV | 0.152 | [0.133, 0.178] | 0.136 | [0.114, 0.161] |
| VFI_20 | Patient | swCV | 0.151 | [0.132, 0.176] | 0.138 | [0.116, 0.163] |
| VFI_21 | Patient | Bak | 0.137 | [0.117, 0.159] | 0.259 | [0.234, 0.283] |
| VFI_22 | Patient | swCV | 0.138 | [0.121, 0.155] | 0.199 | [0.174, 0.221] |
| VFI_23 | Patient | swCV | 0.192 | [0.164, 0.214] | 0.166 | [0.146, 0.188] |
| VFI_24 | Patient | swCV | 0.177 | [0.148, 0.204] | 0.126 | [0.106, 0.145] |
| VFI_25 | Patient | swCV | 0.152 | [0.127, 0.176] | 0.171 | [0.154, 0.194] |
| VFI_26 | Patient | swCV | 0.167 | [0.144, 0.194] | 0.152 | [0.131, 0.170] |
| VFI_27 | Patient | Bak | 0.179 | [0.160, 0.203] | 0.424 | [0.395, 0.453] |

**Supplementary Table S12.** Results from topology weighting by iterative sampling of subtrees (‘*Twisst*’) under three soil reference groups (15 possible topologies). We use one representative isolate per burrow, given the observed clonality. For each isolate, we report the rate at which the dominant topology switched between adjacent windows, and the average run of the dominant topology in units of number of windows and kb. Values denote the mean and 95% confidence interval from the 100 bootstrap replicates.

|  |  | **Switch rate** | | **Runs (in windows)** | | **Runs (in kb)** | |
| --- | --- | --- | --- | --- | --- | --- | --- |
| **Isolate** | **Source** | **Mean** | **95% CI** | **Mean** | **95% CI** | **Mean** | **95% CI** |
| Bakersfield_1  (clonal with Bakersfield_2) | Soil | 0.833 | [0.811, 0.859] | 1.20 | [1.16, 1.23] | 68.3 | [65.9, 70.8] |
| Bakersfield_3 | Soil | 0.829 | [0.803, 0.848] | 1.20 | [1.18, 1.24] | 68.5 | [66.5, 70.7] |
| Bakersfield_4 | Soil | 0.847 | [0.823, 0.874] | 1.18 | [1.14, 1.21] | 67.1 | [64.6, 69.2] |
| Bakersfield_5 | Soil | 0.839 | [0.808, 0.866] | 1.19 | [1.15, 1.23] | 67.5 | [65.3, 70.3] |
| Coalinga_1 | Soil | 0.803 | [0.769, 0.829] | 1.24 | [1.20, 1.29] | 70.7 | [68.3, 73.2] |
| Coalinga_3 | Soil | 0.808 | [0.781, 0.838] | 1.23 | [1.19, 1.27] | 70.4 | [67.7, 73.0] |
| Tracy_1 | Soil | 0.817 | [0.792, 0.844] | 1.22 | [1.18, 1.26] | 69.4 | [67.3, 71.8] |
| Carrizo_1  (clonal with Carrizo_2) | Soil | 0.782 | [0.750, 0.812] | 1.27 | [1.23, 1.33] | 72.6 | [70.0, 76.0] |
| Carrizo_3  (clonal with Carrizo_4 and 5) | Soil | 0.792 | [0.758, 0.822] | 1.26 | [1.21, 1.31] | 71.9 | [69.4, 75.3] |
| McKittrick_1  (clonal with McKittrick_2) | Soil | 0.792 | [0.767, 0.812] | 1.26 | [1.23, 1.30] | 71.9 | [69.6, 74.4] |
| VFI_1 | Patient | 0.815 | [0.786, 0.843] | 1.22 | [1.18, 1.27] | 69.8 | [67.3, 72.5] |
| VFI_2 | Patient | 0.814 | [0.780, 0.845] | 1.23 | [1.18, 1.28] | 70.0 | [67.2, 72.8] |
| VFI_3 | Patient | 0.813 | [0.782, 0.843] | 1.23 | [1.18, 1.27] | 69.9 | [67.3, 72.6] |
| VFI_4 | Patient | 0.838 | [0.808, 0.867] | 1.19 | [1.15, 1.23] | 67.9 | [65.4, 70.4] |
| VFI_5 | Patient | 0.823 | [0.793, 0.861] | 1.21 | [1.16, 1.26] | 68.9 | [65.6, 72.0] |
| VFI_6 | Patient | 0.807 | [0.784, 0.837] | 1.23 | [1.19, 1.27] | 70.4 | [67.7, 72.7] |
| VFI_7 | Patient | 0.819 | [0.790, 0.847] | 1.22 | [1.18, 1.26] | 69.3 | [67.0, 71.8] |
| VFI_8 | Patient | 0.815 | [0.789, 0.843] | 1.22 | [1.18, 1.26] | 69.5 | [66.8, 71.9] |
| VFI_9 | Patient | 0.813 | [0.782, 0.842] | 1.23 | [1.18, 1.27] | 69.9 | [67.3, 72.8] |
| VFI_10 | Patient | 0.821 | [0.790, 0.851] | 1.21 | [1.17, 1.26] | 69.0 | [66.3, 71.6] |
| VFI_11 | Patient | 0.834 | [0.801, 0.863] | 1.20 | [1.16, 1.24] | 68.1 | [65.6, 70.9] |
| VFI_12 | Patient | 0.829 | [0.802, 0.857] | 1.20 | [1.16, 1.24] | 68.5 | [66.0, 70.8] |
| VFI_13 | Patient | 0.834 | [0.808, 0.860] | 1.20 | [1.16, 1.23] | 68.1 | [65.7, 70.7] |
| VFI_14 | Patient | 0.798 | [0.772, 0.829] | 1.25 | [1.20, 1.29] | 71.3 | [68.4, 73.8] |
| VFI_16 | Patient | 0.777 | [0.748, 0.808] | 1.28 | [1.23, 1.33] | 73.2 | [70.5, 76.1] |
| VFI_17 | Patient | 0.825 | [0.796, 0.851] | 1.21 | [1.17, 1.25] | 68.9 | [66.5, 71.8] |
| VFI_18 | Patient | 0.805 | [0.773, 0.833] | 1.24 | [1.20, 1.29] | 70.6 | [68.2, 73.7] |
| VFI_19 | Patient | 0.820 | [0.785, 0.848] | 1.22 | [1.18, 1.27] | 69.5 | [67.0, 72.4] |
| VFI_20 | Patient | 0.820 | [0.787, 0.848] | 1.22 | [1.18, 1.27] | 69.5 | [67.3, 72.2] |
| VFI_21 | Patient | 0.827 | [0.799, 0.857] | 1.21 | [1.16, 1.25] | 68.6 | [65.7, 71.1] |
| VFI_22 | Patient | 0.821 | [0.794, 0.850] | 1.21 | [1.17, 1.25] | 69.3 | [66.5, 71.6] |
| VFI_23 | Patient | 0.803 | [0.774, 0.836] | 1.24 | [1.19, 1.29] | 70.9 | [67.5, 73.4] |
| VFI_24 | Patient | 0.802 | [0.771, 0.829] | 1.24 | [1.20, 1.29] | 71.0 | [68.3, 73.9] |
| VFI_25 | Patient | 0.825 | [0.794, 0.861] | 1.21 | [1.16, 1.26] | 68.8 | [65.6, 71.9] |
| VFI_26 | Patient | 0.798 | [0.774, 0.826] | 1.25 | [1.21, 1.29] | 71.4 | [68.8, 73.7] |
| VFI_27 | Patient | 0.792 | [0.762, 0.830] | 1.26 | [1.20, 1.31] | 71.7 | [68.2, 74.4] |

**Supplementary Table S13.** Source metadata for previously published (“legacy”) *Coccidioides immitis* genomes. Year, location, and source details are based on metadata provided on the Sequence Read Archive (not available for all samples). Here ‘location’ refers to the clinic location or putative exposure location for patient samples. Two-letter state abbreviations are used for states, and ‘SJV’ indicates San Joaquin Valley.

| **Isolate** | **Year** | **Location** | **Source** | **Ref** |
| --- | --- | --- | --- | --- |
| B11019 | 2016 | WA | Patient (testicle) | Monroy-Nieto et al. 2023^24^ |
| B11034 | 2014 | WA (travel to Tucson, AZ) | Patient (lung) | Monroy-Nieto et al. 2023 |
| B11035 | 2015 | WA (or travel) | Patient (cerebrospinal fluid) | Monroy-Nieto et al. 2023 |
| B11057 | 2014 | OR | Patient (lung) | Monroy-Nieto et al. 2023 |
| B11198 | 2015 | WA (travel to CA & Mexico) | Patient (tissue) | Monroy-Nieto et al. 2023 |
| B11517 | 2016 | WA (travel to CA) | Patient (throat) | Monroy-Nieto et al. 2023 |
| B12219 | 2016 | WI (travel to Southwestern U.S.) | Patient (lung wash) | Monroy-Nieto et al. 2023 |
| B12220 | 2016 | OR (travel to CA) | Patient (sputum) | Monroy-Nieto et al. 2023 |
| B12495 | 2016 | WA (travel to CA) | Patient (Cerebrospinal fluid) | Monroy-Nieto et al. 2023 |
| B12496 | 2016 | WA (travel to CA & Mexico) | Knee wound | Monroy-Nieto et al. 2023 |
| B12526 | 2016 | OR (travel to AZ and CA) | Patient (lung wash) | Monroy-Nieto et al. 2023 |
| B13956 | 2017 | WA (potential travel to CA) | Patient | Monroy-Nieto et al. 2023 |
| B14131 | 2017 | OR | Patient | Monroy-Nieto et al. 2023 |
| B14132 | 2017 | OR (travel to CA) | Patient (lymph node) | Monroy-Nieto et al. 2023 |
| B14133 | 2017 | OR | Patient (tissue) | Monroy-Nieto et al. 2023 |
| B14135 | 2017 | OR (travel to CA) | Patient (lung wash) | Monroy-Nieto et al. 2023 |
| B14286 | 2017 | WA (travel to TX) | Patient (wound) | Monroy-Nieto et al. 2023 |
| B14288 | 2017 | OR | Patient | Monroy-Nieto et al. 2023 |
| B14298 | 2018 | WA (non-recent travel to CA) | Patient (wound) | Monroy-Nieto et al. 2023 |
| B15142 | 2018 | OR | Patient | Monroy-Nieto et al. 2023 |
| B15145 | 2018 | OR | Patient | Monroy-Nieto et al. 2023 |
| B15146 | 2018 | OR | Patient | Monroy-Nieto et al. 2023 |
| B15257 | 2018 | WA | Patient (back abscess) | Monroy-Nieto et al. 2023 |
| B15368 | 2018 | WA | Patient (bronchial wash) | Monroy-Nieto et al. 2023 |
| B16338 | NA | Mexico | Patient (bronchial wash) | Monroy-Nieto et al. 2023 |
| B16339 | 2018 | OR | Patient (body fluid) | Monroy-Nieto et al. 2023 |
| B16364 | 2018 | Mexico | Patient (pleural fluid) | Monroy-Nieto et al. 2023 |
| B16536 | 2018 | OR | Patient | Monroy-Nieto et al. 2023 |
| B16692 | 2018 | OR | Patient | Monroy-Nieto et al. 2023 |
| B17554 | 2018 | WA | Patient | Monroy-Nieto et al. 2023 |
| B17567 | NA | WA | Patient | Monroy-Nieto et al. 2023 |
| B17635 | 2019 | WA | Patient | Monroy-Nieto et al. 2023 |
| Coahuila_1 | NA | Mexico | Patient | Engelthaler et al. 2016^25^ |
| San_Diego_1 | NA | San Diego, CA | Patient | Engelthaler et al. 2016 |
| SJV_10 | NA | SJV, CA | Patient | Engelthaler et al. 2016 |
| SJV_11 | NA | SJV, CA | Patient | Engelthaler et al. 2016 |
| SJV_2 | NA | SJV, CA | Patient | Engelthaler et al. 2016 |
| SJV_4 | NA | SJV, CA | Patient | Engelthaler et al. 2016 |
| SJV_5 | NA | SJV, CA | Patient | Engelthaler et al. 2016 |
| SJV_7 | NA | SJV, CA | Patient | Engelthaler et al. 2016 |
| SJV_8 | NA | SJV, CA | Patient | Engelthaler et al. 2016 |
| SJV_9 | NA | SJV, CA | Patient | Engelthaler et al. 2016 |
| UCLA_293 | NA | Southern CA | Patient | No associated publication |
| Guerrero_1 | NA | South Mexico | Patient | Engelthaler et al. 2016 |
